# Vibrotactile stimulation on the upper back requires stronger vibration during goal-directed reaching but does not impede reaching performance

**DOI:** 10.64898/2026.10.13.755060

**Authors:** Alice E. Atkin, Bernard Marius ’t Hart, Cari M. Whyne, Sebastian Tomescu, Bradley H. Strauss, Qingguo Li, Denise Y.P. Henriques

## Abstract

Vibrotactile stimulation is a promising method of biofeedback, with potential applications for ergonomic posture and balance, rehabilitation, and motor learning. These applications require balancing tactile perception with motor performance and cognitive workload. In this study, participants performed a detection task, a reaching task, and a concurrent reach-and-detect task, with vibrotactile signals delivered on the hand or upper back. Sensitivity to stimuli, motor performance, and cognitive workload were compared for each task. Sensitivity was suppressed during movement, but this suppression was greater on hand compared to the upper back. Movement time was slowed by vibration on the hand but not the upper back. Mental workload and task complexity increased for the reach-and-detect task compared to the individual tasks. We conclude that vibrotactile feedback devices worn on the upper back do not impede motor performance, with only minor tactile suppression. This could be overcome by presenting highly salient or personalized feedback.

**Practitioner Summary:** This study evaluated the use of a wearable vibrotactile feedback device during a concurrent reaching task. Concurrent reaching reduced feedback detection, but this reduction was smaller when the upper back was vibrated compared to the hand, with less disruption of reaching performance. Concurrent reaching and detection was judged to be more mentally demanding than either task alone. Vibrotactile feedback devices can be used effectively on the upper back, but users should be aware of mental fatigue.

## Background

Vibrotactile stimulation is a promising method of biofeedback, with potential applications for ergonomic posture and balance (Ballardini et al., 2020; Bonin et al., 2026; Kelly et al., 2023), rehabilitation (Bae & Park, 2023; Bonab et al., 2026; Moesgen et al., 2022), and motor learning (Islam & Lim, 2022; Risi et al., 2019; Shah et al., 2022). Successful use of vibrotactile feedback devices requires people to balance tactile perception with motor performance and cognitive workload. These demands vary depending on their applications, however. During movement, people’s sensitivity to vibrotactile stimulation changes. Perception of tactile stimuli is weaker on moving parts of the body during goal-directed movements, a phenomenon called “tactile gating” or “tactile suppression” (Arikan et al., 2024; Chapman et al., 1987; Juravle et al., 2017; Kilteni & Ehrsson, 2022; Williams et al., 1998). Tactile suppression may limit the utility of vibrotactile feedback devices in applied settings by reducing people’s sensitivity to feedback as they work. One solution to this problem may be to wear the vibrating motors on a relatively distal and immobile part of the body, such as the back, since suppression effects are proximate to the moving effector (Colino et al., 2014; Williams et al., 1998). Tactile suppression is also enhanced during conditions of increased cognitive workload, which may further reduce sensitivity to feedback (McManus et al., 2023). Conversely, monitoring for vibrotactile signals carries a cognitive workload and increases stress (DeLucia & Greenlee, 2022), which may distract from efficient motor task performance. The effective use of vibrotactile feedback devices therefore depends not just on perceiving the vibration, but also on whether processing that information changes motor performance or cognitive workload.

Tactile suppression typically follows a distinct time course, beginning prior to movement, peaking early during movement, and then recovering towards the end of movement (Buckingham et al., 2010; Tatai et al., 2026; Vitello et al., 2012). This typical time course corresponds to body state estimates, which are low before movement and then increase throughout the movement (Fraser & Fiehler, 2018). When uncertainty is low, the sensorimotor system is not reliant on sensory feedback, and more tactile information can be suppressed without consequence. From this, researchers have concluded that tactile suppression is dynamically weighted according to how important sensory feedback is for properly executing a planned movement (Voudouris & Fiehler, 2017; 2021), or according to task demands (McManus et al. 2023).

One family of well-characterized motor task is the Fitts Aiming task. In Fitts Aiming, people perform reaching movements to targets of varying width and distance using their hand, a stylus, a computer mouse, or some other type of input device (Fitts, 1954; MacKenzie, 2018). Movement time (MT) typically increases according to an index of difficulty (ID), which can be defined according to the width of a target and the distance that must be traversed to reach it. This is because movements to smaller or further targets require greater reach precision, which is achieved by producing slower movements. MT and ID also allow researchers to calculate information throughput (TP) which represents the limit of performance for a given type of movement (Gori et al., 2018; MacKenzie, 2018; Thorne, 2006; Zhai, 2004). Fitts’-style reaching tasks are useful because they offer a tidy way of measuring how motor performance changes with task difficulty and cognitive workload, with predictable hypotheses.

Performing two tasks simultaneously can reduce performance on one or both tasks due to cognitive load limits (see, ex. Morrison et al., 2015). The effects of workload on performance can be manipulated and measured using a dual-task paradigm. In such a paradigm, two tasks are performed separately to provide a baseline measure of performance, and then also performed concurrently. The increased difficulty of performing two tasks concurrently can result in reduced performance on one or both tasks and an increase in workload. The cognitive load of a task can be assessed subjectively using questionnaires such as the NASA Task Load Index (NASA-TLX; Hart, 2006; Hart & Staveland, 1988) or modified versions such as the Surgical Task Load Index (SURG-TLX; Wilson et al., 2011). These questionnaires measure a task’s subjective mental demands, as well as other relevant performance factors such as physical and temporal demands, task complexity, stress and distraction. If monitoring for vibrotactile signals during motor tasks results in reduced performance or high workload this could render vibrotactile feedback systems ineffective, burdensome, or unsafe for use within clinical or occupational settings, where many tasks are cognitively demanding and errors can have serious consequences (Dias et al., 2018).

In a previous study from our group, we found that use of a vibrotactile feedback device can improve people’s posture during a fine motor task performed with the hands, reducing their time spent in risky postures (Bonin et al., 2026). Participants performed a modified hand tool dexterity task while responding to vibrotactile feedback about their posture. This study extends that work by simultaneously characterizing tactile perception, motor performance, and workload, rather than focusing on postural correction. Our vibrotactile feedback device is intended to be worn on the back, which is less sensitive to tactile stimulation than other areas of the body, such as the hand, but this location may nevertheless be advantageous if it proves less susceptible to suppression effects. Use of the feedback device in the previous study didn’t increase people’s cognitive workload compared to a control condition, but it’s possible that increasing people’s workload by increasing task difficulty will inhibit people’s ability to process the vibrotactile stimuli. Finally, the motor task in that study only provided us with a single measure of task completion time, with no ability to perform more fine-grained analyses involving trial-by-trial movement time, variability, velocity, path trajectories, or throughput. The current study was designed to address these outstanding questions.

The goal of this study was to characterize how vibrotactile stimuli delivered during goal-directed reaching influences tactile perception, motor performance, and subjective workload. We hypothesized that sensitivity to vibrotactile stimuli would be lower during a reach-and-detect task compared to a solo detection task, requiring stronger vibration in order to be detectable. We also hypothesized that tactile sensitivity would be lower on the upper back than the hand, but that suppression effects would be larger on the hand than the back. Movement time (MT) on the reaching task was hypothesized to be slower in a reach-and-detect task compared to a solo reaching task, but we did not expect the location of the vibrotactile motors to affect MT or any other measure of motor performance. These comparisons allowed us to assess whether the upper back is a practical location for delivering vibrotactile signals during reaching while minimizing interference with motor performance.

## Methods

### Participants

40 participants were initially recruited for this study. 6 participants were excluded because they did not complete both study sessions. 4 participants were excluded because they exhibited a low sensitivity for tactile detection, as evidenced through flat slopes on their psychometric curve functions. One participant was excluded because they reported detecting vibration on every trial of the reach-and-detect task. Thus, the final sample for analysis consisted of 29 participants (24 women, 4 men, 1 non-binary, age range = 18-23, mean age = 18.97 ± 1.20, right-handed = 27).

This study received ethical approval from the Office of Research Ethics at York University in Toronto, Canada, project #766, and was conducted in accordance with the Declaration of Helsinki. All participants provided written consent for their participation and received course credit as compensation. Participants consented to have their anonymized data made publicly available on OSF as a condition of their participation: https://osf.io/n8rqz/overview

### Equipment

Vibrotactile stimuli were delivered using a custom-made vibrotactile stimulation device, which consisted of an Arduino NANO 33 IoT connected to two 5V eccentric rotating mass (ERM) vibration motors (Adafruit). Motors were affixed with medical tape to either the upper back ∼50mm to the left and right of the T3 vertebra, or to the index finger of the dominant hand, on the dorsal and ventral surfaces of the intermediate phalange. Responses to vibrational stimuli were recorded using a computer keyboard.

The reaching task was performed using a 430.4 x 287mm Intuos Pro digitizing tablet and stylus (Wacom Technology Corporation). Stimuli were presented on a 19-inch monitor (ViewSonic Corporation) and were responded to with a keyboard.

### Stimuli and Procedure

In this study, participants performed a detection task, a reaching task, and a reach-and-detect task which combined them (Figure 1). The study was divided into two sessions at least 24 hours apart, to limit fatigue from repetitive reaching. In Session 1, participants sequentially performed the detection task, the reaching task, the reach-and-detect task, and the reaching task again. In Session 2, participants performed the detection task and the reach-and-detect task. Participants wore the motors on the back or the hand in separate sessions, with the order counterbalanced across participants.

**Figure 1.**
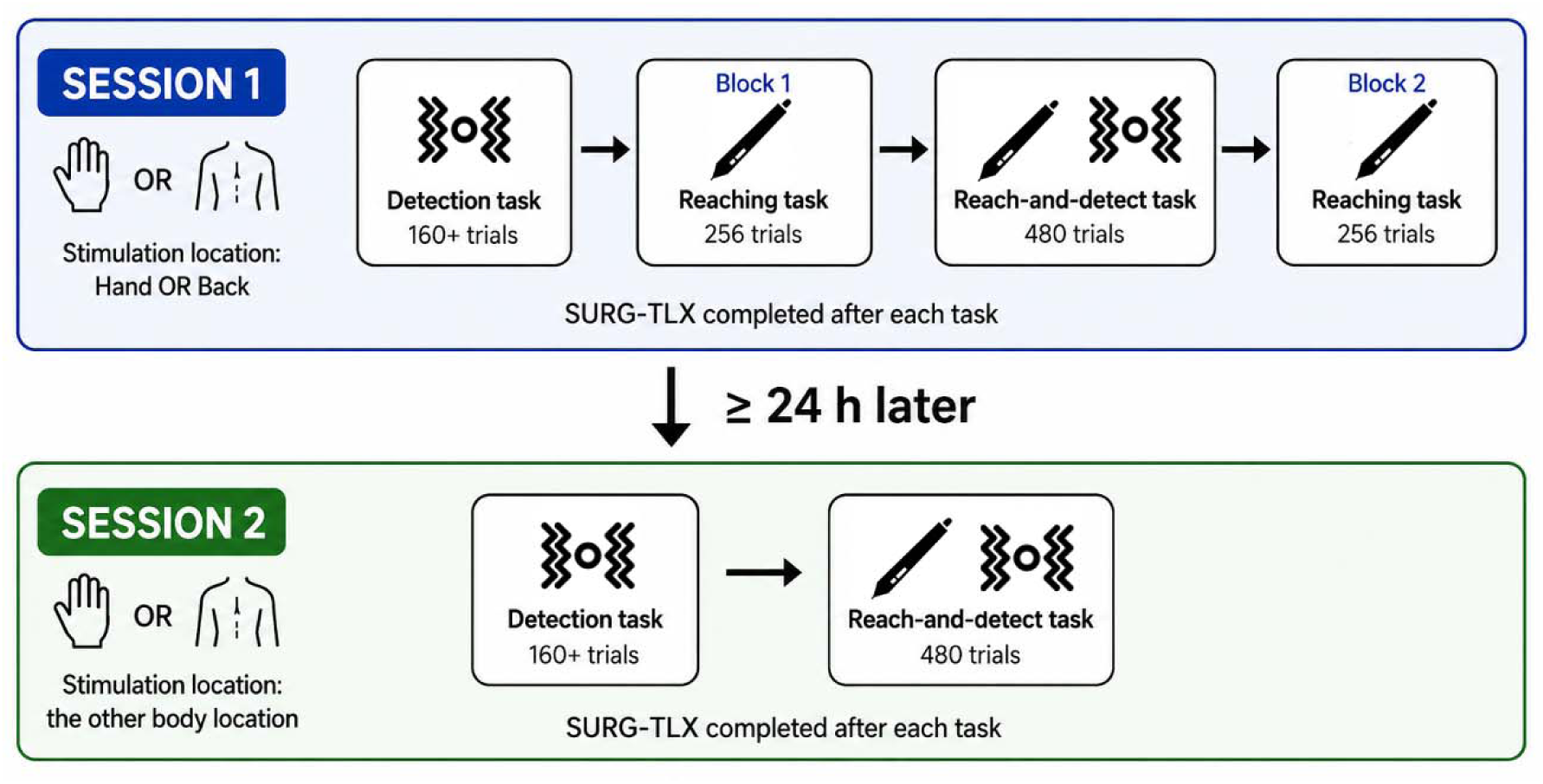
Study Design. In Session 1, participants performed, in sequence, the detection task, the reaching task, the reach-and-detect task, and then the reaching task again. In Session 2, participants performed the detection and reach-and-detect tasks. Participants who wore the vibrotactile motors on the hand in Session 1 wore the motors on the back in Session 2, and vice versa. Trial numbers for each task are per-participant.

#### Detection Task

For the detection task, a staircasing procedure was used to present the stimuli. Four separate staircases were presented in an interspersed manner, with two staircases starting at the maximum vibration strength (127 arbitrary units) and two staircases starting at the minimum vibration (1 a.u.). Within each staircase, vibrational intensity was decreased when the previous stimulus was detected, and increased when the previous stimulus was not detected. Step size was initially set at 24 units, with step sizes decreasing by 3 units following each reversal. Each staircase ran until a minimum of 40 trials had been presented, with at least 12 reversals in stimulus intensity. This means the detection task consisted of a minimum of 160 trials per participant. Stimulus duration was 50ms for all trials, which is common for tactile suppression tasks (see, ex. Arikan et al., 2024; Buckingham et al., 2010; or McManus et al., 2023). Pilot testing showed that a 50ms duration produced good psychometric curves across a range of intensities relative to shorter or longer durations, with their respectively lower and higher detection rates.

The task began with 8 practice trials so that participants could familiarize to the task. 4 practice trials had vibration, presented at the maximum intensity of 127 a.u. for 200ms duration to ensure detection. The other 4 practice trials had no vibration. Participants responded by pressing the Right Arrow key if they detected vibration, and the Left Arrow key if they did not. Feedback about correct or incorrect responses was provided for the practice trials only, and participants were reminded about the response button mapping if they responded incorrectly.

On the monitor, trial onset was cued via the presentation of a red dot (5mm radius) against a black background. The vibrational stimulus was presented at a random timepoint between 500-1500ms following trial onset. The red dot turned blue 1500ms after trial onset to cue the response. The blue dot stayed onscreen until a keypress was provided. The inter-trial interval between participant response and the next trial onset was 500ms.

The detection task was performed twice and had two location conditions, with the vibrotactile motors worn on the hand and the upper back in separate sessions.

#### Reaching Task

For the reaching task, stimuli consisted of a red dot (2.5mm radius) representing the stylus cursor, a circular start position, and a circular aiming target. Stimuli were presented against a black background. The start position was a 5mm white circle, while the aiming target was either a 5mm circle (Small Target) or 10mm circle (Large Target). The aiming target was white until the participant entered it with their cursor and held that position for 500ms, at which point it turned yellow to cue a keyboard response. The location of the aiming target on one trial became the start position for the next trial.

There were 4 target positions, located in the upper and lower left and right corners of the screen (Figure 2). On each trial, the target always appeared in one of the 2 locations on the opposite side of the screen from the start position, ensuring that participants performed reciprocal leftward and rightward aiming movements across the tablet from trial to trial. The distance between the centre of the start position and the centre of the aiming target was either 20cm (Near Distance) or 22.36cm (Far Distance). Each combination of target sizes and distances was presented 64 times across two blocks, for a total of 512 trials. Participants always pressed the Right Arrow key when the target turned yellow to advance to the next trial, as this task had no vibrational stimuli to detect or not detect. Immediately following response, the target transformed into the start position for the next trial. Participants had to hold their cursor within the start position for 500ms before the next target appeared. Participants received three breaks during each block for a self-determined length of time.

**Figure 2.**
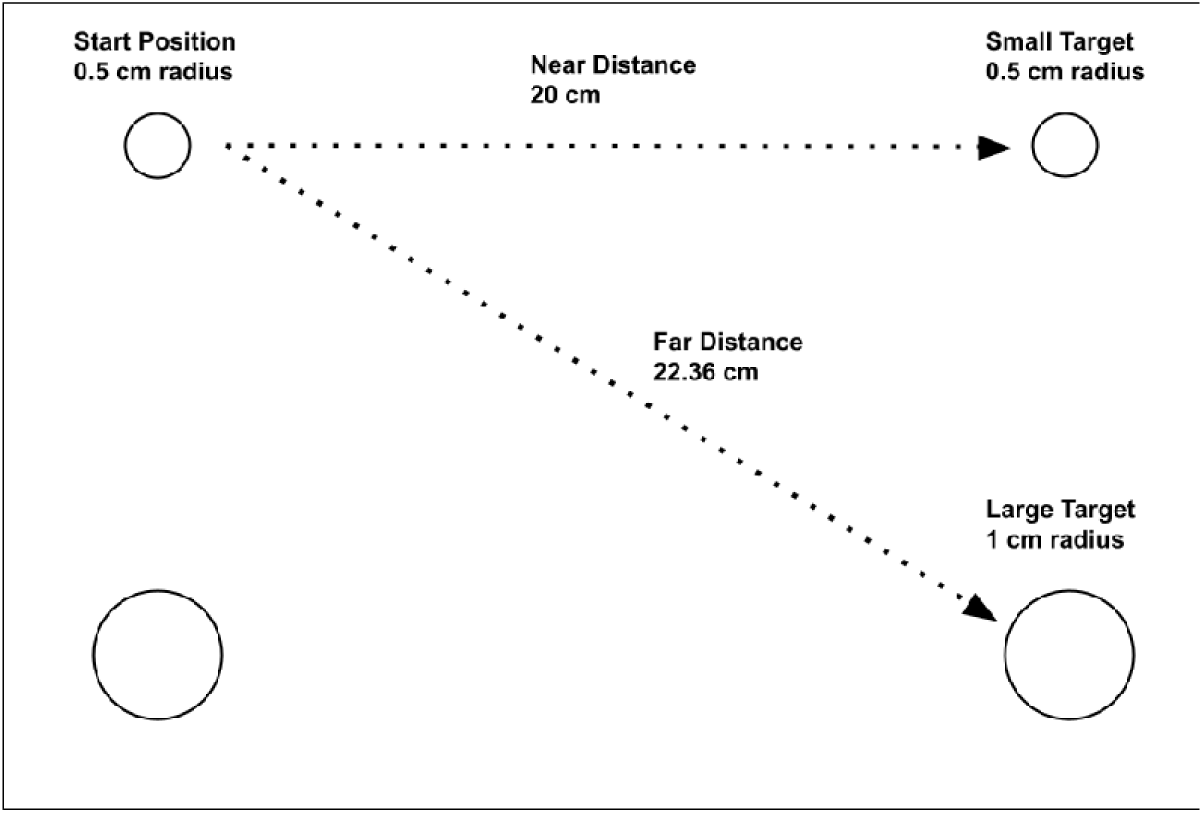
A representation of target sizes and distances in the reaching and the reach-and-detect tasks (not to scale). Targets could be located in the top-left, top-right, bottom-left, or bottom-right of the computer screen. Targets were either small (0.5cm) or large (1cm), and distances were either near (20cm) or far (22.361cm). Target position on one trial became the starting position for the next trial. Targets were always on the opposite side of the screen from the starting position. Only the start position and current-trial target position were visible to participants during each trial; the two other, possible target positions were not visible.

Two bouts of the reaching task were performed in Session 1, before and after the reach-and-detect task. Participants were assigned to cohorts depending on whether they wore the vibrotactile motors on the hand or the upper back during Session 1, and cohorts were compared to determine if wearing the motors on the hand caused physical interference with reaching.

#### Reach-and-Detect Task

The reach-and-detect task combined elements of the detection task and the reaching task. Vibration was presented for 50ms at 5 set intensities (0, 30, 60, 90, and 120 units). Each vibrational intensity was presented 96 times in pseudo-random order, for a total of 480 trials. Trials were also divided into three timing conditions. Vibration was either presented immediately after the onset of the reach target, before movement onset had occurred (*Pre-Onset condition*); immediately after movement onset, when the cursor position exited the start position (*At Onset condition*); or in the middle of the movement, 500ms after the cursor left the start position (*During Movement condition*). Each timing condition was presented 32 times at each vibrational intensity, for a total of 160 trials per timing condition. Visual stimuli on the monitor were identical to the reaching task. Each combination of target sizes and distances was presented 120 times, for a total of 480 trials. Similar to the detection task, participants responded to each trial by pressing the Right Arrow key if they felt a vibration, or the Left Arrow key if they did not.

As with the detection task, the reach-and-detect task was performed in two bouts, with the vibrotactile motors worn on the hand and the upper back in separate sessions.

### SURG-TLX Questionnaire

Immediately following each task, participants completed the Surgery Task Load Index (SURG-TLX) questionnaire (Wilson et al., 2010). The SURG-TLX provides a subjective, retrospective measure of cognitive workload during a task on six dimensions: mental demands, physical demands, temporal demands, task complexity, situational stress, and distraction. The SURG-TLX was administered using Google Forms (Google, Inc.) and participants rated each dimension on a 20-point Likert scale. Participants were instructed to rate the task they had just completed, rather than the study as a whole.

### Data Analysis

Data processing was performed in Python (v. 3.13, Python Software Foundation). Statistical analysis was performed in R (v. 4.4.2, The R Foundation for Statistical Computing), using the packages “afex” (Singmann et al., 2025), “emmeans” (Lenth et al., 2026), “effectsize” (Ben-Shachar et al., 2026), and “tidyverse” (Wickham, 2023). For each within-subjects ANOVA analysis, Greenhouse-Geiser sphericity correction and Holm correction for multiple comparisons were applied when appropriate.

#### Vibrotactile Detection

50% detection thresholds were calculated by fitting a probit regression model with margins to individual participant data as illustrated in Figure 3. The probit model produced a psychometric curve representing the relationship between vibration strength and the probability that participants detected the vibration, thus allowing us to determine the intensity at which vibration is detected 50% of the time. Estimating detection thresholds from psychometric functions provides a more robust measure of tactile sensitivity than detection accuracy at a single intensity because it incorporates responses across a range of strengths. A probit model was appropriate because it assumes normally-distributed data, thus giving the model a conceptual link to signal detection theory and our d’ prime sensitivity analysis (DeCarlo, 1998; Horowitz & Savin, 2001). The model was characterized by up to four parameters. The mean and standard deviation of the normal distribution of responses were mandatory parameters, while lapse rate and guess rate were optional parameters. Lapse rate represents the possibility that participants do not detect even high-strength vibration due to inattention, while guess rate represents the possibility that some detection reports for low-strength stimuli represent guessing rather than valid detection. 50% detection thresholds were compared with a two-way within-subjects ANOVA, using the factors “Task” (Detection x Reach-and-Detect and “Location” (Back x Hand).

**Figure 3.**
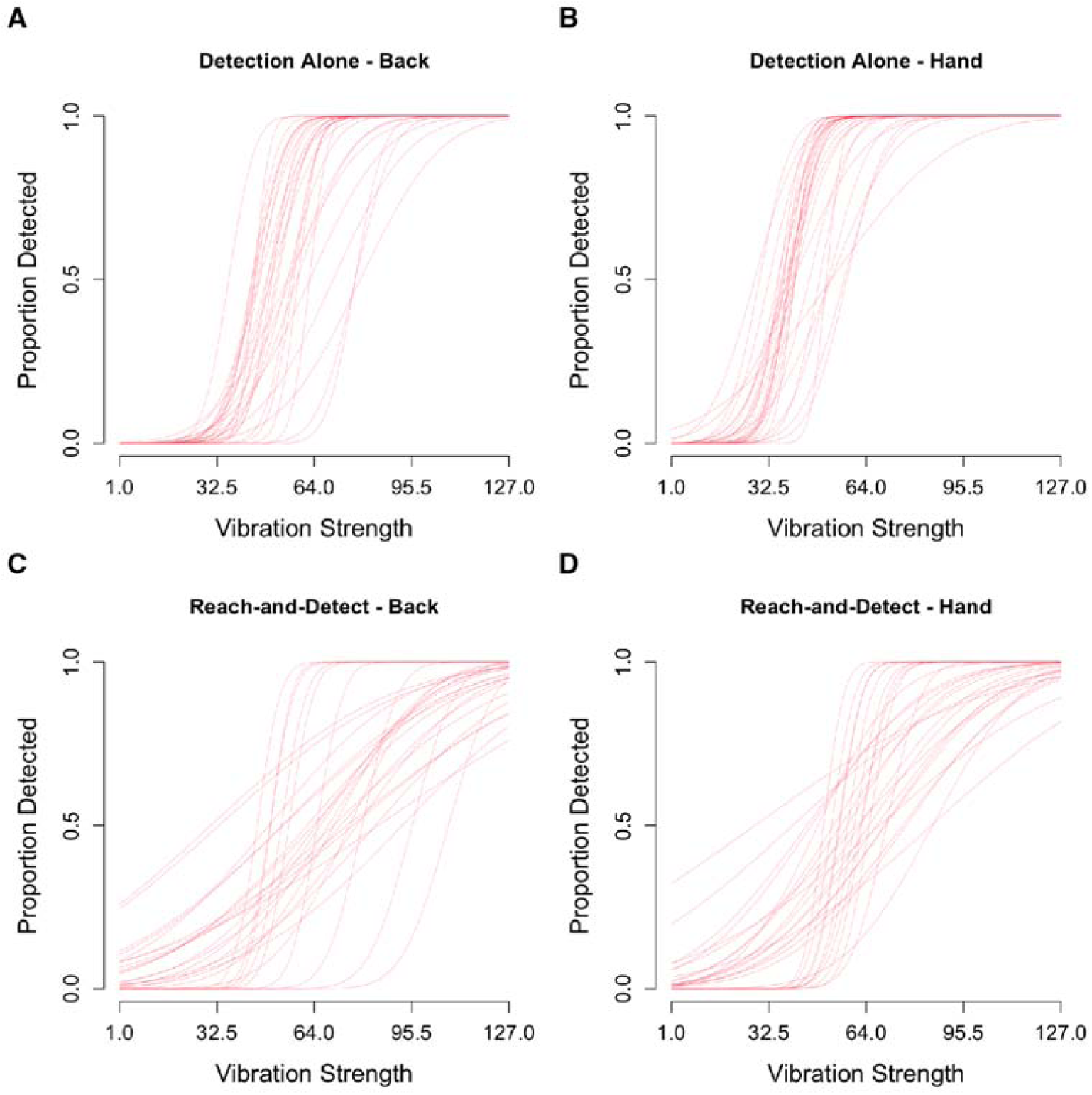
Psychometric curves for each participant (N=29), representing the relationship between the strength of vibration and the proportion of trials on which the vibration was detected. A: Psychometric curves when vibration was delivered to the back during the Detection task. B: Psychometric curves when vibration was delivered to the hand during the Detection task. C: Psychometric curves when vibration was delivered to the back during the Reach-and-Detect task. D: Psychometric curves when vibration was delivered to the hand during the Reach-and-Detect task.

We also measured sensitivity another way for the reach-and-detect task, using d’. In signal detection theory, d’ represents the standardized difference between the mean of the signal and noise distributions, and is calculated as each participant’s hit rate minus their false alarm rate (Stanislaw & Todorov, 1999):

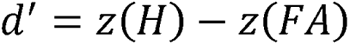

The 1 / 2N correction from Macmillan & Kaplan (1985) was applied to extreme hit or false alarm rates of 0 or 1. d’ was only measured for the reach-and-detect task because the detection task used a staircasing procedure with vibration on every trial, meaning there were no true false alarms, only hits or misses. 20% of reach-and-detect trials, by contrast, had no vibration, so a d’ analysis is possible. We compared d’ for vibration on the back compared to the hand, and for each vibration timing condition.

A suppression effect was calculated by subtracting each participant’s 50% detection threshold during the detection task from their 50% detection thresholds at each vibration timing during the reach-and-detect task. Because sensitivity differs between the back and the hand, suppression scores were calculated relative each participants’ detection thresholds for each location. This subtraction produced a suppression score, with positive values indicating suppressed sensitivity to vibration during the reach-and-detect task, and negative values representing enhanced sensitivity. Suppression scores were compared using a two-way within subjects ANOVA, using the factors “Location” (Back x Hand) and “Vibration Timing” (Pre-Onset x At Onset x During Movement).

#### Motor Performance

Movement time (MT), movement variability, path length, and throughput were analyzed for the reaching task and the reach-and-detect task using within-subjects ANOVA’s, with the exception of the cohort analysis for the reaching task, which used a mixed ANOVA with cohort as the between-subjects factor. These results are reported in the Results section below. We also analyzed peak velocity, time to peak velocity, average velocity, and acceleration, and this information is reported in a supplementary R notebook: https://osf.io/n8rqz/overview

MT was measured for the phase of movement when the cursor had left the start position, but had not yet entered the target. Movement variability was defined as the standard deviation of the signed error in the y-dimension upon reaching the target’s near-side circumference in the x-dimension. Trials had their coordinate frames rotated so that the x and y coordinates were aligned on all trials, regardless of the location of the start point and target, with the x-dimension representing the straight-line distance from start position to target, and the y-dimension being perpendicular to that line. Path length was calculated for the same phase of movement as MT, and represents the distance travelled by the cursor. To calculate throughput, an index of difficulty (ID) was calculated for each trial using the Shannon formulation for Fitts’ Law (MacKenzie, 2018):

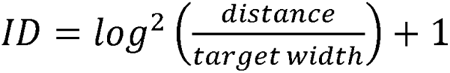

Throughput was then calculated as the ratio between ID and MT:

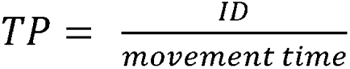

Throughput was calculated because it integrates movement time and precision into a single measure, allowing us to analyze whether detecting vibrotactile stimuli on the back or the hand affected overall motor performance. TP was compared across each value of ID and task (*Reaching, Reach-and-Detect* and *Reach-and-Detect [Hand]*) using a two-way within-subjects ANOVA. All other measures were compared using a three-way within subjects ANOVA, with the factors being “Task”, “Target Size” (*small* and *large*) and “Distance” (*near* and *far*).

#### SURG-TLX

Responses to each of the six SURG-TLX dimensions were analysed using one-way within-subjects ANOVAs, using task as the factor (*Detection, Reaching*, and *Reach-and-Detect*).

#### Data Exclusions

Trials during the reaching and reach-and-detect tasks were excluded from analysis if MT exceeded 2500ms, which we took as an indication that participants had responded carelessly on that trial. This resulted in approximately 0.9% of all trials being excluded from analysis (*Reaching:* 0.5%; *Reach-and-Detect:* 0.7%; *Reach-and-Detect [Hand]:* 1.5%). Inspection of MT’s across the reaching task revealed a practice effect within the first block, with MT progressively decreasing before stabilizing. To ensure that performance on this task reflected stable performance and was comparable to the reach-and-detect task, only the second half of the first reaching block (128 of 256 trials) was retained for analysis. These trials were merged with the data from the second reaching block into a single reaching task. The first 12 trials from the second reaching block were also discarded as warmup trials. Thus, the final analysis of the reaching task included 372 trials per participant, minus additional exclusions for excessive MT.

## Data Availability

Participant data and supplementary materials are publicly available on OSF: https://osf.io/n8rqz/overview

Experiment, data processing and analysis scripts are available on GitHub: https://github.com/thealiceatkin/Vibrotactile-stimulation-during-goal-directed-reaching.git

## Results

### Tactile Perception

To characterize how vibrotactile detection during goal-directed reaching influenced tactile perception, we first examined detection thresholds and movement-related suppression before evaluating motor performance and subjective workload.

### Detection Thresholds

As illustrated in Fig. 4A, participants required stronger vibration to detect tactile stimuli while reaching than during the detection task (*Detection:* 47.88 a.u. ± 11.28; *Reach-and-Detect:* 63.37 a.u. ± 15.59). Detection thresholds were higher when vibration was delivered to the upper back (*red;* 60.29 a.u. ± 15.78) than to the hand (*blue*; 50.96 a.u. ± 14.11), with the difference being greater during reaching than for the detection task, *F* [1, 28] = 69.358, *p* < .001, η^2^_p_ = .712; *F* [1, 28] = 13.830, *p* < .001, η^2^_p_ = 0.331; interaction: *F* [1, 28] = 8.407, *p* = .007, η^2^_p_ = 0.231). The slope of the psychometric function was shallower during reaching (16.24 a.u. ± 15.96) compared to the detection task (9.73 ± 4.82), indicating reduced tactile sensitivity during movement, *F* [1, 28] = 58.192, *p* < .001, η^2^_p_ = .675.

**Figure 4.**
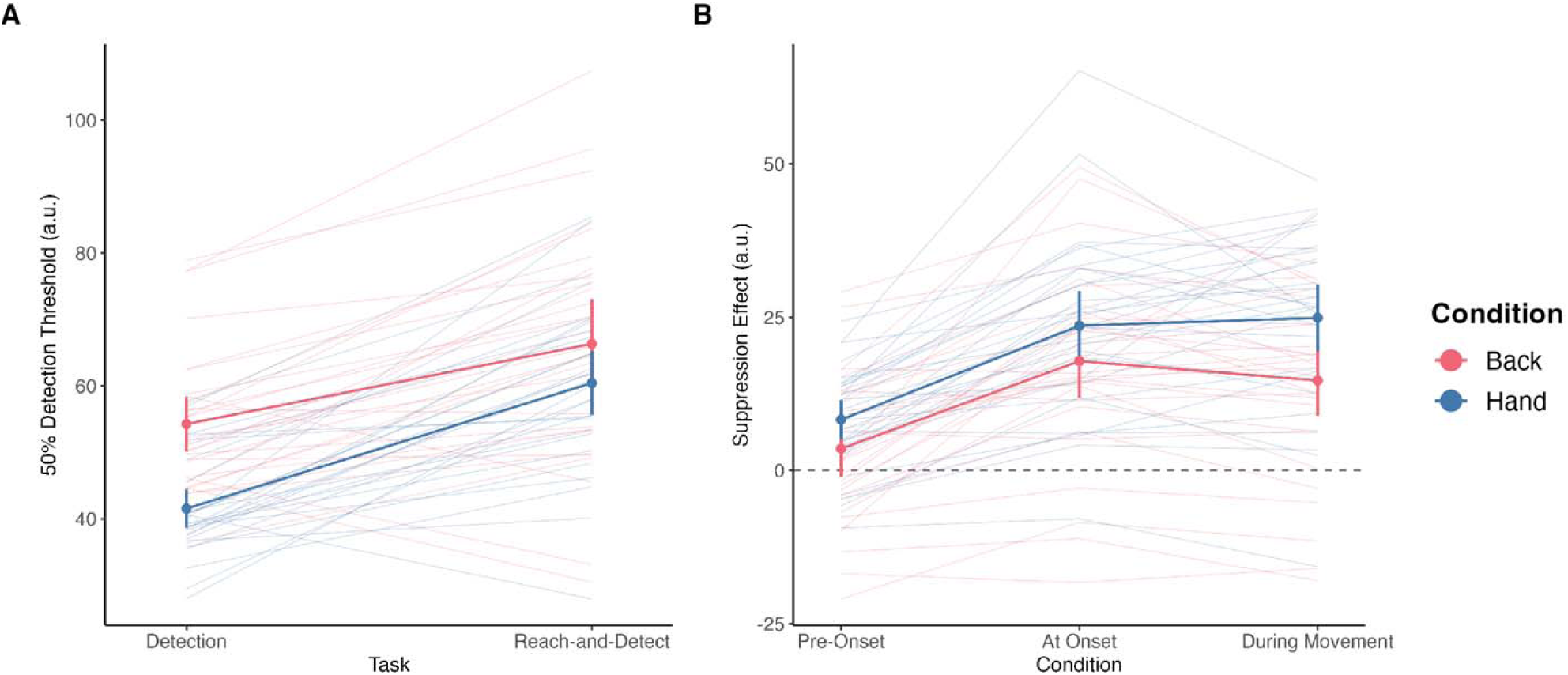
A: 50% Detection thresholds depending on task and the location of vibration. Detection threshold represents the vibrational intensity required for participants to detect a vibration 50% of the time. The detection threshold is higher in the reach-and-detect compared to the detection task, and higher for the back compared to the hand. B: The suppression effect represents the difference in 50% detection thresholds for the reach-and-detect task compared to the detection task, and has positive values when the threshold is higher for the reach-and-detect task. The suppression effect was greater for the hand compared to the back, and greater when vibration was delivered at the onset of movement or during movement compared to pre-movement.

Figure 3 presents the psychometric curves for the back and the hand in the detection task and the reach-and detect task, illustrating the rightward shift in the detection threshold for the back (3A; 60.29 a.u. ± 15.78) relative to the hand (3B; 50.96 a.u. ± 14.11) and the shallower slopes in the reach-and-detect task (3C and 3D; *Detection:* 9.73 a.u. ± 4.82; *Reach-and-Detect:* 26.23 a.u. ± 15.96).

### Suppression Effect

Suppression was greater for vibration delivered at movement onset or during movement rather than before movement, and greater for the hand than the back (Fig. 4B). Suppression was calculated as the difference between the reach-and-detect and detection task thresholds. As illustrated in Figure 4B, this effect differed depending upon the location of vibration during the reach-and-detect task, *F* [1, 28] = 7.289, *p* <=.012, η^2^_p_ = .207, and the timing of vibration, *F* [1.94, 54.42] = 49.481, *p* < .001, η^2^_p_ = .639. There was also an interaction between the location and timing of vibration, *F* [1.77, 49.21] = 4.703, *p* = .007, η^2^_p_ = .144. The suppression effect was larger for vibration on the hand compared to the back (*Back [red]:* 12.02 a.u. ± 15.49; *Hand [blue]:* 18.93 a.u. ± 14.86). Post-hoc tests revealed that the suppression effect was larger when the vibration was delivered at movement onset or during movement, compared to when the vibration was delivered prior to movement onset, *t* [28] = 9.184, *p* < .001 (*Pre-Onset:* 5.91 a.u. ± 10.51; *At Onset:* 20.72 a.u. ± 15.49; *During Movement:* 19.79 a.u. ± 15.53). The At Onset and During Movement conditions were collapsed for this post-hoc analysis because they did not differ from each other, *t* [28] = 0.615, *p* = .544.

Sensitivity (d’) was analyzed for the reach-and-detect task only. Sensitivity was reduced for vibration on the hand and reduced for vibration delivered at onset or during movement. There was also an interaction between location and vibration timing. These findings were supported by the analysis, which showed significant effects of location, *F* [1, 28] = 17.916, *p* < .001, η^2^_p_ = .390, timing of vibration, *F* [1.97, 55.16] = 16.351, *p* < .001, η^2^_p_ = .369, and an interaction, *F* [1.94, 54.31] = 4.936, *p* =.011, η^2^_p_ = .150. Thus, sensitivity was significantly lower on the hand than the upper back, however this difference was greater when vibration was delivered before movement onset and smaller when vibration was delivered at onset or during movement (*Pre-Onset:* back = 1.69 ± 0.46, hand = 2.12 ± 0.47; *At Onset:* back = 1.48 ± 0.46, hand = 1.78 ± 0.54; *During Movement:* back = 1.49 ± 0.42, hand = 1.62 ± 0.49).

### Motor Performance

Movement times were longer when the hand (*blue*) was vibrated during the reach-and-detect task than when the upper back (*red*) was vibrated or during the reaching task (*black*), (Fig. 5A). Movement time also increased systematically with ID across all three tasks, with nearly parallel functions indicating a similar effect of task difficulty regardless of vibration location. These observations of MT were supported by a significant effect of task, *F* [1.57, 43.84] = 10.545, *p* < .001, η^2^_p_ = .274. Post-hoc tests showed that average MT was longer in the reach-and-detect task when the hand (*blue*) was vibrated, compared to the reaching task (*black*), *t* [28] = 3.573, *p* = .004, and when the upper back (*red*) was so vibrated, *t* [28] = 3.973, *p* = .002 (*Reaching:* 1.03s ± 0.20; *Back:* 1.05s ± 0.19; *Hand:* 1.13s ± 0.23). The effect of ID was similar across all tasks, *F* [2.07, 57.94] = 418.723, *p* < .001, η^2^_p_ = .937. No interaction was observed.

**Figure 5.**
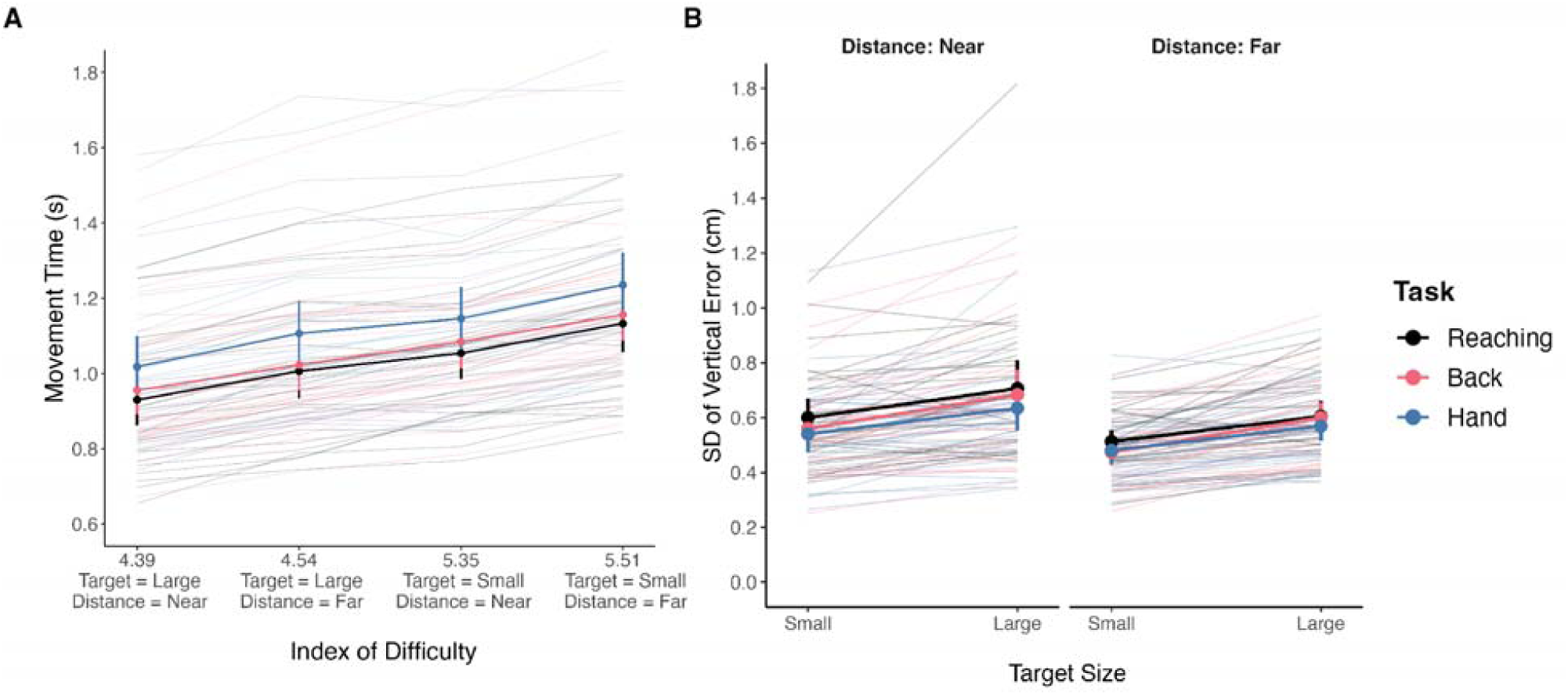
Mean movement time and SD of vertical error. A: MT depending on task and Index of Difficulty. MT was longer for trials with a higher level of difficulty, and was longer in the reach-and-detect task when the hand was vibrated. B: SD of vertical error when the movement reached the target, depending on task, target size, and distance. Reach error variability was the same for all tasks, but more precise for far distances and small targets.

To rule out the possibility that wearing the motors on the hand physically interfered with movement, we divided our participants into two cohorts, those who performed the reaching task with the motors on the upper back, and those who performed the reaching task with the motors on the hand. We then performed a mixed ANOVA, with cohort as the between-subjects factor and ID as the within-subjects factor. The results showed no effect of cohort *F* [1, 27] = 0.206, *p* = .653, η^2^_p_ = .008, and no interaction with ID, *F* [2.38, 64.21] = 0.957, *p* = .402, η^2^_p_ = .034. We therefore find no evidence that wearing the motors on the hand resulted in slower movements during the reaching task, when there was no vibration.

### Movement Variability

Participants were equally precise in their reaching movements regardless of task (Fig. 5B). The analysis showed an effect of target size, *F* [1, 28] = 121.519, *p* < .001, η^2^_p_ = .813, and distance, *F* [1, 28] = 21.258, *p* < .001, η^2^_p_ = .432, but task was not significant, *F* [1, 28] = 2.016, *p* = .144, η^2^_p_ = .067. Reach error variability was reduced for smaller targets, as one might expect, and also for far targets.

### Path Length

As to be expected, path length depended primarily on the distance to the target from the starting position, *F* [1, 28] = 25791.161, *p* < .001, η^2^_p_ = .999. Reaches to further targets (21.57 cm ± 0.48) were longer than those for nearer targets (Near: 19.27 cm ± 0.51). Reaches to larger targets (20.10 cm ± 1.21) followed slightly shorter paths than for smaller targets (20.73 cm ± 1.22), *F* [1, 28] = 2237.058, *p* < .001, η^2^_p_ = .988. There was no main effect of task and no interactions were observed.

### Throughput

Throughput was lower when the hand was vibrated compared to the back or the reaching task. The TP analysis showed main effects of both task, *F* [1.55, 43.35] = 10.667, *p* < .001, η^2^_p_ = .276, and index of difficulty, *F* [2.24, 62.84] = 135.825, *p* < .001, η^2^_p_ = .829. There was no interaction between the factors. Vibration on the hand during the reaching task significantly reduced TP (*Reaching*: 5.25 bits/s ±0.96; *Back*: 5.08 bits/s ± 0.83; *Hand*: 4.80 bits/s ± 0.87). Post-hoc tests showed that TP was higher for the reaching task compared to the reach-and-detect task when vibration was on the hand, *t* [28] = 3.725, *p* = .003, but not when vibration was on the back, *t* [28] = 1.792, *p* = .084. Between the two locations, TP was higher when vibration was on the back compared to the hand, *t* [28] = 3.748, *p* = .003. TP varied with index of difficulty, but not linearly as might be expected (*4.39*: 4.98 bits/s ± 0.92; *4.55*: 4.75 bits/s ± 0.85; *5.36*: 5.34 bits/s ± 0.91; *5.51*: 5.10 bits/s ± 0.84). This pattern indicates that TP was higher for small targets (ID = 5.36 or 5.51) compared to large targets (ID = 4.39 or 4.55), but was also higher for far targets (ID = 4.39 or 5.36) compared to near targets (ID = 4.55 or 5.51). Thus, the aspect of difficulty that drove increased TP in this study was target size, not distance.

### Cognitive Workload

#### Mental Demands

In this study, the reach-and-detect task was more mentally demanding than the detection or reaching tasks (Fig. 6A). The reaching task was also more demanding than the detection task. The SURG-TLX analysis showed a significant effect of task, *F* [1.94, 54.41] = 35.949, *p* < .001, η^2^_p_ = .562. Post-hoc tests showed that the detection task differed from the reaching task, *t* [28] = 3.389, *p* = .002, and from the reach-and-detect task, *t* [28] = 8.666, *p* < .001. The reaching task also differed from reach-and-detect, *t* [28] = 5.105, p < .001. The reaching task was more mentally demanding than the detection task, and the reach-and-detect task was more demanding than either of the other tasks (*Detection:* 5.62 ± 2.38; *Reaching:* 7.85 ± 3.69; *Reach-and-Detect:* 10.76 ± 3.39).

**Figure 6.**
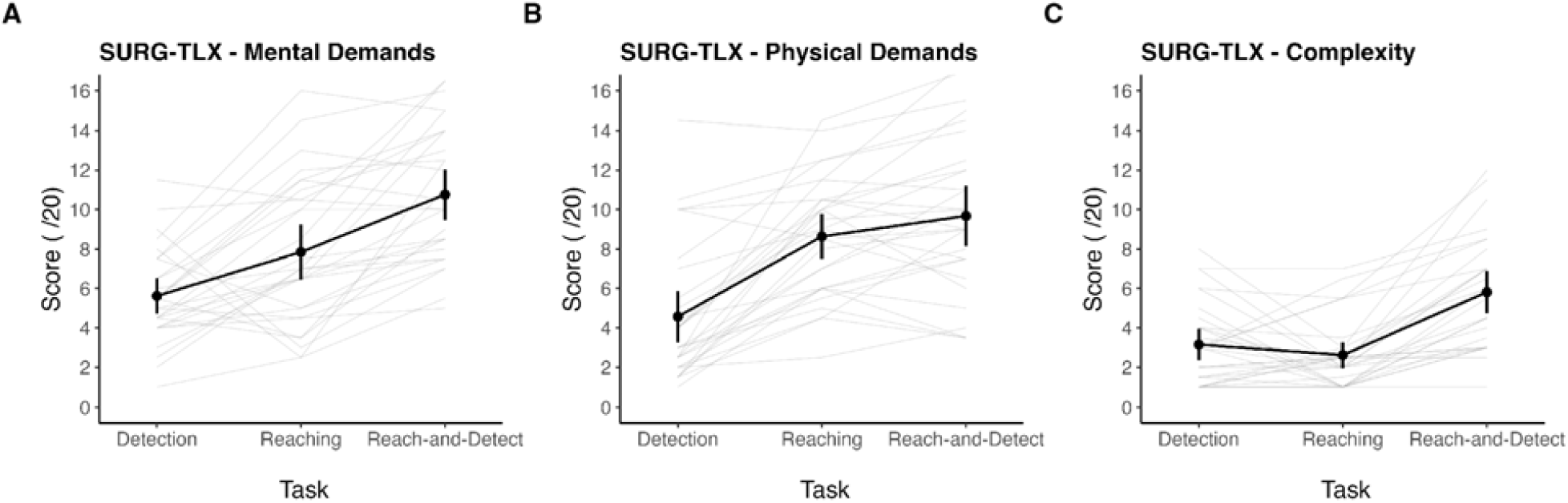
Mean SURG-TLX responses for the questions on A) mental demands, B) physical demands, and C) task complexity, depending on the task. Participants found the reaching task more mentally and physically demanding than the detection task, but not more complex. The reach-and-detect task was judged to be more mentally demanding and more complex than either the detection or reaching tasks, and was more physically demanding than the detection task but not the reaching task. Back and hand locations for the detection task and the reach-and-detect task were collapsed together for these analyses.

#### Physical Demands

The reaching and reach-and-detect tasks were more physically demanding than detection (Fig. 6B). The SURG-TLX analysis showed a significant effect of task, *F* [1.59, 44.46] = 44.605, *p* < .001, η^2^_p_ = .614. Post-hoc tests showed that detection differed from reaching, *t* [28] = 7.350, *p* < .001, and from reach-and-detect, *t* [28] = 7.372, *p* < .001. The reaching task also differed from reach-and-detect, *t* [28] = 2.353, *p* = .026. The reaching task was more physically demanding than the detection task, and the reach-and-detect task was more demanding than either of the other tasks (*Detection:* 4.57 ± 3.45; *Reaching:* 8.64 ± 3.02; *Reach-and-Detect:* 9.67 ± 4.02).

#### Temporal Demands

For the temporal demands question, participants showed no effect of task, *F* [1.42, 39.71] = 2.224, *p* = .135, η^2^_p_ = .074. Temporal demands did not differ between the detection and reaching tasks, or between the reach-and-detect task and the other tasks (*Detection:* 5.67 ± 3.42; *Reaching:* 4.38 ± 2.56; *Reach-and-Detect:* 4.97 ± 2.39).

#### Task Complexity

The reach-and-detect task was judged to be more complex than the detection or reaching tasks (Fig. 6C). Participants showed a significant effect of task, *F* [1.95, 54.72] = 30.850, *p* < .001, η^2^_p_ = .524. Post-hoc tests showed that the detection task differed from the reach-and-detect task, *t* [28] = 5.686, *p* < .001, and the reaching task also differed from the reach-and-detect task, *t* [28] = 7.609, *p* < .001. Detection did not differ from reaching, *t* [28] = 1.283, *p* = .210. Participants found the reach-and-detect task to be more complex than the other tasks (*Detection:* 3.16 ± 2.08; *Reaching:* 2.62 ± 1.74; *Reach-and-Detect:* 5.81 ± 2.83).

#### Stress

For the stress question, participants showed no effect of task, *F* [1.90, 53.26] = 2.363, *p* = .106, η^2^_p_ = .078. Perceived stress did not differ between the detection and reaching tasks, or between the reach-and-detect task and the other tasks (*Detection:* 3.83 ± 2.72; *Reaching:* 3.57 ± 2.91; *Reach-and-Detect:* 4.41 ± 3.47).

#### Distraction

For the distraction question, participants showed a significant effect of task, *F* [1.71, 47.91] = 3.536, *p* = .044, η^2^_p_ = .112. However, post-hoc tests did not reveal significant differences between any two tasks. Detection did not differ from reaching, *t* [28] = 0.380, *p* = .707, or from reach-and-detect, *t* [28] = 1.861, *p* = .146, and reaching also did not differ from reach-and-detect, *t* [28] = 2.366, *p* = .075 (*Detection:* 2.81 ± 2.33; *Reaching:* 2.71 ± 2.54; *Reach-and-Detect:* 3.55 ± 3.01).

## Discussion

In this study, we combined a vibrotactile detection task with a Fitts-style reaching task and compared performance on each task individually against performance when they were performed concurrently. The vibrotactile motors were attached to participants’ dominant hand or their upper back, and performance was compared across these two locations. As hypothesized, the results showed tactile suppression during the reach-and-detect task, with lower sensitivity to vibrotactile stimulation at movement onset or during movement, but not prior to movement (Fig. 4B). This suppression effect was larger for vibration on the hand compared to the upper back, although a suppression effect on the back was still evident. The upper back was less sensitive to vibration, also as expected, with the mean detection threshold occurring at a higher vibrational intensity than for the hand (Fig. 4A). Wearing the vibrotactile motors on the hand also notably affected motor performance during the reach-and-detect task by slowing movement times (Fig. 5A). Performing the two tasks concurrently was subjectively perceived as being more mentally demanding and more complex than performing either task by itself (Fig. 6A and 6C).

Tactile suppression was observed during movement when the vibrotactile motors were worn on the upper back, although it was not as large as for the hand (Fig. 4B). This finding was unexpected, as suppression is typically localized proximate to the effector (Colino et al., 2014; Colino & Binsted, 2016; Williams et al., 1998). In Colino et al. (2014), suppression just prior to a reaching and grasping movement was reliably found only on the forearm of the moving arm, and not on the fingers or anywhere on the contralateral limb. Williams et al. (1998) found a strong relationship between proximity to the effector, in this case the index finger, and the magnitude of suppression. In that study, participants were almost half as likely to detect stimuli on the fingers or hand as they were stimuli on the shoulders or pectoral muscles, the closest analogues to our upper back location. One possible explanation for the strong upper back suppression observed in our study is that our side-to-side reaching movements activated the muscles of the shoulder and upper back. Van Hulle et al. (2013) found tactile suppression for vibrotactile stimuli delivered to the back during back movements. In that study, however, participants reached for a target by bending over a table, which involves the upper back and shoulders much more substantially than our task. Although it was possible for our participants to perform the reaching movements using minimal shoulder involvement, relying only on a rotation of the elbow and wrist, most participants likely did use their shoulder to some extent. If so, this suggests that wearing vibrotactile feedback devices on the upper back does not entirely avoid the issue of tactile suppression, at least for tasks that involve the whole arm, rather than merely the hand or fingers. Nevertheless, wearing vibrating motors on the upper back reduces suppression effects compared to wearing them on the hand.

Another possible explanation is that the tactile suppression observed during movement for both locations is partly due to workload. During reaching, participants must attend to their movement trajectory and use somatosensory feedback to make corrections so their hand acquires the target (Fiehler et al., 2019; Voudouris & Fiehler, 2021). The more critical somatosensory feedback is to performance, such as when reaching for somatosensory rather than visual targets or when vision is occluded, the more suppression is typically observed (Colino et al., 2017; Gertz et al., 2017; Voudouris & Fiehler, 2017). One study by McManus et al. (2023) found a correlation between higher subjective task load and increased suppression during reaching, which they attribute to a prioritization of reach-relevant signals under conditions of high load. Consistent with this finding, the reach-and-detect task was rated as more mentally and physically demanding and more complex than the detection task (Fig. 6). Participants in our study were not instructed to prioritize either the detection or the reaching aspects of the Reach-and-Detect task, but if high task demands induced them to prioritize the reaching task over detection then this could produce suppression. Thus, vibrotactile stimulation during motor tasks may be limited in utility when workload is high or tasks are complex as this may reduce sensitivity to vibrotactile signals (DeLucia & Greenlee, 2023).

The upper back was less sensitive to vibrotactile stimulation than the hand, requiring stronger vibration to reach a detection threshold (Fig. 4A). This threshold was well below the maximum intensity of our vibrotactile device, however, and in applied settings it would be possible to apply strong, supra-threshold vibration in all circumstances, as was done in Bonin et al. (2026), rather than varying the intensity as in this study. Vibrotactile feedback could also be personalized for individual users depending upon their sensitivity (Luzhnica et al., 2017). Thus, there appears to be little risk that wearing vibrotactile feedback devices on the upper back will have limited utility due to lower sensitivity relative to other bodily locations. Consistent with our previous study, we find that users can reliably detect strong vibration on the upper back even while performing a simultaneous motor task (Bonin et al., 2026). These findings suggest that the upper back is a viable location for delivering vibrotactile feedback, so long as the vibration is clearly suprathreshold.

During the reach-and-detect task, wearing the vibrotactile motors on the upper back generally did not affect reaching performance compared to when reaching was performed without vibration, with no changes to the speed or variability of movement (Fig. 5). The one exception to this finding was slower average velocity during the reach-and-detect task, which is reported in the supplementary materials. Since average velocity was slower for vibration on the hand as well, this result could be a general effect of performing two tasks concurrently, and the increased task complexity and mental workload involved. It could be due to fatigue from repetitive reaching, as our participants rated the tasks involving reaching as being much more physically demanding than the detection task (Fig. 6B; Dupuis et al., 2021; Takahashi et al., 2006). Overall, vibrating the upper back during movement had minimal impact on reaching, especially when compared to the hand. We therefore find no strong evidence that monitoring for vibrotactile stimulation on the upper back during manual tasks will impede task performance. This finding supports further evaluation of the upper back as a location for task-relevant vibrotactile feedback.

Slower MT and lower throughput when the hand was vibrated may have occurred because either the vibration itself or the suppression effect interfered with the execution of movement. Tactile suppression requires movements that surpass a critical speed, approximately 5 cm/s, which our participants greatly exceeded (Cybulska-Klosowicz et al., 2011). Above this threshold, evidence is mixed as to whether suppression increases or decreases with velocity (Angel & Malenka, 1982; Fraser & Fiehler, 2018; Gertz et al., 2017; Schmidt et al., 1990), but suppression is usually greatest during the early stage of movement, when velocity peaks (Buckingham et al., 2010; Tatai et al., 2026; Vitello et al., 2012). Our results could suggest a negative relationship between speed and suppression, or it could be the case that participants deliberately slowed their movements to counter strong suppression and improve their ability to detect the vibration. The current study cannot dissociate these two hypotheses. Overall, we find that delivering vibration on the hand reduces speed of movement compared to no vibration, whereas providing vibration on the upper back does not. These findings suggest that the upper back is a suitable location for delivering vibrotactile signals during manual activities that involve reaching.

## Limitations

Several aspects of this study limit our ability to draw conclusions from our results and generalize them to other studies. The study was performed in two sessions, and several participants were excluded from the final analysis because they did not return for the second session. Although we cannot determine why these participants dropped out of the study, attrition may have reduced the representation of participants who found the procedure difficult or uncomfortable. Because of this attrition, our results should be generalized cautiously to applied settings, where users are likely to exhibit the full range of capabilities and tolerances for using any similar vibrotactile feedback system. Our study included a limited number of target sizes and locations, and the magnitude of our observed effects may differ for targets that require different speeds, precision, force, or whole arm movement. Our motor results should therefore be generalized cautiously to other targets and tasks (Zhai et al., 2004). Finally, our participants were all young adults between the ages of 18-23. The narrow age range of our sample may limit generalization across the broader working-age population.

## Conclusion

In this study, we tested people’s ability to detect vibrotactile signals while performing a concurrent goal-directed reaching task. The findings identify perceptual, motor, and workload constraints that should be considered when designing vibrotactile feedback systems for use during manual activity. Vibrotactile stimulation has been proposed as an affordable and reliable method of providing feedback about posture, balance, and other physiological factors in occupational or rehabilitation settings, or during motor learning (Bae & Park, 2023; Ballardini et al., 2020; Bonab et al., 2026; Bonin et al., 2026; Islam & Lim, 2022; Kelly et al., 2023; Moesgen et al., 2022; Risi et al., 2019; Shah et al., 2022). To be effective for these purposes, however, vibration must be detectable without impeding performance on concurrent tasks. The results showed that tactile suppression was greater at the time of movement onset or during movement for vibration on the hand compared to the upper back, while movement time was slowed for the hand but not the upper back. We therefore find little evidence that wearing a vibrotactile feedback device on the upper back will substantially impede performance on motor tasks, and only minor evidence that performing a concurrent motor task will impede vibrotactile feedback detection. However, performing vibrotactile detection concurrently with a motor task may increase the perceived complexity of the task, increasing mental fatigue. In applied settings, moderate tactile suppression on the upper back could be overcome by presenting highly salient feedback (Bonin et al., 2026) or by personalizing the device to each user’s sensitivity level (Luzhnica et al., 2017).

## Supporting information

Supplementary Analyses

## Author Contributions Statement

**Alice Elizabeth Atkin** contributed conceptualization, methodology, software, validation, formal analysis, investigation, data curation, writing (original draft), writing (review & editing), visualization, and project administration. **Bernard Mariust’ Hart** contributed conceptualization, methodology, software, formal analysis, data curation, and writing (review & editing). **Cari M. Whyne** contributed conceptualization, methodology, writing (review & editing), supervision, and funding acquisition. **Sebastian Tomescu** contributed conceptualization, methodology, writing (review & editing), supervision, and funding acquisition. **Bradley H. Strauss** contributed conceptualization, methodology, writing (review & editing), supervision, and funding acquisition. **Qingguo Li** contributed conceptualization, methodology, resources, writing (review & editing), supervision, and funding acquisition. **Denise Y.P. Henriques** contributed conceptualization, methodology, validation, formal analysis, resources, writing (original draft), writing (review & editing), visualization, supervision, project administration, and funding acquisition

## Acknowledgements

The authors wish to thank Christopher Giverin at York University for his technical assistance. The authors acknowledge that Claude (v. 4.6, Anthropic, PBC) was consulted for software coding assistance. All code was reviewed and verified for accuracy.

## References

Angel, R. W., & Malenka, R. C. (1982). Velocity-dependent suppression of cutaneous sensitivity during movement. Experimental Neurology, 77(2), 266–274. 10.1016/0014-4886(82)90244-8

Arikan, B. E., Voudouris, D., Straube, B., & Fiehler, K. (2024). Distinct role of central predictive mechanisms in tactile suppression. iScience, 27(8), 110582. 10.1016/j.isci.2024.110582

Bae, S., & Park, H.-S. (2023). Development of immersive virtual reality-based hand rehabilitation system using a gesture-controlled rhythm game with vibrotactile feedback: An fNIRS pilot study. IEEE Transactions on Neural Systems and Rehabilitation Engineering, 31, 3732–3743. 10.1109/TNSRE.2023.3312336

Ballardini, G., Florio, V., Canessa, A., Carlini, G., Morasso, P., & Casadio, M. (2020). Vibrotactile feedback for improving standing balance. Frontiers in Bioengineering and Biotechnology, 8(94). 10.3389/fbioe.2020.00094

Ben-Shachar, M. S., Makowski, D., Lüdecke, D., Patil, I., Wiernik, B. M., Thériault, R., Kelley, K., Stanley, D., Caldwell, A., Burnett, J., Karreth, J., & Waggoner, P. (2026). effectsize: Indices of effect size (Version 1.0.2) [Computer software]. https://cran.r-project.org/web/packages/effectsize/index.html

Bonab, A. K., Camardella, C., Serra, F., Frisoli, A., Posteraro, F., & Leonardis, D. (2026). Leveraging wearable haptics for guidance in virtual rehabilitation: Effects on motor control from an immersive VR setting. Scientific Reports, 16(1), 5513. 10.1038/s41598-026-35092-6

Bonin, W. R., Brost, S., Atkin, A. E., Henriques, D. Y. P., Tomescu, S., Strauss, B. H., Whyne, C. M., & Li, Q. (2026). Vibrotactile feedback using exposure variation analysis to reduce musculoskeletal risk in surgical simulation. Ergonomics, 1–11. 10.1080/00140139.2026.2615116

Buckingham, G., Carey, D. P., Colino, F. L., deGrosbois, J., & Binsted, G. (2010). Gating of vibrotactile detection during visually guided bimanual reaches. Experimental Brain Research, 201(3), 411–419. 10.1007/s00221-009-2050-8

Chapman, C. E., Bushnell, M. C., Miron, D., Duncan, G. H., & Lund, J. P. (1987). Sensory perception during movement in man. Experimental Brain Research, 68(3). 10.1007/BF00249795

Colino, F. L., & Binsted, G. (2016). Time course of tactile gating in a reach-to-grasp and lift task. Journal of Motor Behavior, 48(5), 390–400. 10.1080/00222895.2015.1113917

Colino, F. L., Buckingham, G., Cheng, D. T., Van Donkelaar, P., & Binsted, G. (2014). Tactile gating in a reaching and grasping task. Physiological Reports, 2(3), e00267. 10.1002/phy2.267

Colino, F. L., Lee, J.-H., & Binsted, G. (2017). Availability of vision and tactile gating: Vision enhances tactile sensitivity. Experimental Brain Research, 235(1), 341–348. 10.1007/s00221-016-4785-3

Cybulska-Klosowicz, A., Meftah, E.-M., Raby, M., Lemieux, M.-L., & Chapman, C. E. (2011). A critical speed for gating of tactile detection during voluntary movement. Experimental Brain Research, 210(2), 291–301. 10.1007/s00221-011-2632-0

DeCarlo, L. T. (1998). Signal detection theory and generalized linear models. Psychological Methods, 3(2), 186–205. 10.1037/1082-989X.3.2.186

DeLucia, P. R., & Greenlee, E. T. (2022). Tactile vigilance Is stressful and demanding. Human Factors, 64(4), 732–745. 10.1177/0018720820965294

Dias, R. D., Conboy, H. M., Gabany, J. M., Clarke, L. A., Osterwei, L. J., Avrunin, G. S., Arney, D., Goldman, J. M., Riccardi, G., Yule, S. J., & Zenati, M. A. (2018). Development of an interactive dashboard to analyze cognitive workload of surgical teams during complex procedural care. 2018 IEEE Conference on Cognitive and Computational Aspects of Situation Management (CogSIMA), 77–82. 10.1109/COGSIMA.2018.8423995

Dupuis, F., Sole, G., Wassinger, C., Bielmann, M., Bouyer, L. J., & Roy, J.-S. (2021). Fatigue, induced via repetitive upper-limb motor tasks, influences trunk and shoulder kinematics during an upper limb reaching task in a virtual reality environment. PLOS ONE, 16(4), e0249403. 10.1371/journal.pone.0249403

Fiehler, K., Brenner, E., & Spering, M. (2019). Prediction in goal-directed action. Journal of Vision, 19(9), 10. 10.1167/19.9.10

Fitts, P. M. (1954). The information capacity of the human motor system in controlling the amplitude of movement. Journal of Experimental Psychology, 47(6), 381–391. 10.1037/h0055392

Fraser, L. E., & Fiehler, K. (2018). Predicted reach consequences drive time course of tactile suppression. Behavioural Brain Research, 350, 54–64. 10.1016/j.bbr.2018.05.010

Gertz, H., Voudouris, D., & Fiehler, K. (2017). Reach-relevant somatosensory signals modulate tactile suppression. Journal of Neurophysiology, 117(6), 2262–2268. 10.1152/jn.00052.2017

Gori, J., Rioul, O., & Guiard, Y. (2018). Speed-accuracy tradeoff: a formal information-theoretic transmission scheme (FITTS). ACM Transactions on Computer-Human Interaction, 25(5), 1–33. 10.1145/3231595

Hart, S. G. (2006). Nasa-Task Load Index (NASA-TLX); 20 years later. Proceedings of the Human Factors and Ergonomics Society Annual Meeting, 50(9), 904–908. 10.1177/154193120605000909

Hart, S. G., & Staveland. (1988). Development of NASA-TLX (Task Load Index): Results of empirical and theoretical research. Advances in Psychology, 52, 139–183. 10.1016/S0166-4115(08)62386-9

Horowitz, J. L., & Savin, N. E. (2001). Binary Response Models: Logits, Probits and Semiparametrics. Journal of Economic Perspectives, 15(4), 43–56. 10.1257/jep.15.4.43

Islam, M. S., & Lim, S. (2022). Vibrotactile feedback in virtual motor learning: A systematic review. Applied Ergonomics, 101, 103694. 10.1016/j.apergo.2022.103694

Juravle, G., Binsted, G., & Spence, C. (2017). Tactile suppression in goal-directed movement. Psychonomic Bulletin & Review, 24(4), 1060–1076. 10.3758/s13423-016-1203-6

Kelly, N. A., Althubaiti, A., Katapadi, A. D., Smith, A. G., Nyirjesy, S. C., Yu, J. H., Onwuka, A. J., & Chiang, T. (2023). Association of vibrotactile biofeedback with reduced ergonomic risk for surgeons during tonsillectomy. JAMA Otolaryngology–Head & Neck Surgery, 149(5), 397. 10.1001/jamaoto.2023.0044

Kilteni, K., & Ehrsson, H. H. (2022). Predictive attenuation of touch and tactile gating are distinct perceptual phenomena. iScience, 25(4), 104077. 10.1016/j.isci.2022.104077

Lenth, R. V., Piaskowski, J., Banfai, B., Bolker, B., Buerkner, P., Giné-Vázquez, I., Hervé, M., Jung, M., Love, J., Miguez, F., Riebl, H., & Singmann, H. (2026). emmeans: Estimated marginal means, aka least-squares means (Version 2.0.3) [Computer software]. https://cran.r-project.org/web/packages/emmeans/index.html

Luzhnica, G., Stein, S., Veas, E., Pammer, V., Williamson, J., & Smith, R. M. (2017). Personalising vibrotactile displays through perceptual sensitivity adjustment. Proceedings of the 2017 ACM International Symposium on Wearable Computers, 66–73. 10.1145/3123021.3123029

MacKenzie, I. S. (2018). Fitts’ Law. In K. L. Norman & J. Kirakowski (Eds.), The Wiley Handbook of Human Computer Interaction (1st ed., pp. 347–370). Wiley. 10.1002/9781118976005.ch17

Macmillan, N. A., & Kaplan, H. L. (1985). Detection theory analysis of group data: Estimating sensitivity from average hit and false-alarm rates. Psychological Bulletin, 98(1), 185–199. 10.1037/0033-2909.98.1.185

McManus, M., Schütz, I., Voudouris, D., & Fiehler, K. (2023). How visuomotor predictability and task demands affect tactile sensitivity on a moving limb during object interaction in a virtual environment. Royal Society Open Science, 10(12), 231259. 10.1098/rsos.231259

Moesgen, T., Salovaara, A., Pouta, E., Pyykko, R., & Xiao, Y. (2022). Vibrotactile motion guidance for stroke rehabilitation: A comparative study. 2022 6th International Conference on Medical and Health Informatics, 147–152. 10.1145/3545729.3545759

Morrison, N. M. V., Burnham, D., & Morrison, B. W. (2015). Cognitive load in cross-modal dual-task processing. Applied Cognitive Psychology, 29(3), 436–444. 10.1002/acp.3122

Risi, N., Shah, V., Mrotek, L. A., Casadio, M., & Scheidt, R. A. (2019). Supplemental vibrotactile feedback of real-time limb position enhances precision of goal-directed reaching. Journal of Neurophysiology, 122(1), 22–38. 10.1152/jn.00337.2018

Schmidt, R. F., Schady, W. J. L., & Torebjörk, H. E. (1990). Gating of tactile input from the hand: I. Effects of finger movement. Experimental Brain Research, 79(1), 97–102. 10.1007/BF00228877

Shah, V. A., Thomas, A., Mrotek, L. A., Casadio, M., & Scheidt, R. A. (2023). Extended training improves the accuracy and efficiency of goal-directed reaching guided by supplemental kinesthetic vibrotactile feedback. Experimental Brain Research, 241(2), 479–493. 10.1007/s00221-022-06533-1

Singmann, H., Bolker, B., Westfall, J., Aust, F., Ben-Shachar, M. S., Højsgaard, S., Fox, J., Lawrence, M. A., Mertens, U., Love, J., Lenth, R., & Christensen, R. H. B. (2025). afex: Analysis of factorial experiments (Version 1.5-1) [Computer software]. https://cran.r-project.org/web/packages/afex/index.html

Stanislaw, H., & Todorov, N. (1999). Calculation of signal detection theory measures. Behavior Research Methods, Instruments, & Computers, 31(1), 137–149. 10.3758/BF03207704

Takahashi, C. D., Nemet, D., Rose-Gottron, C. M., Larson, J. K., Cooper, D. M., & Reinkensmeyer, D. J. (2006). Effect of muscle fatigue on internal model formation and retention during reaching with the arm. Journal of Applied Physiology, 100(2), 695–706. 10.1152/japplphysiol.00140.2005

Tatai, F., Voudouris, D., Straub, D., Fiehler, K., & Rothkopf, C. A. (2026). Tactile suppression during movement as optimal integration of somatosensory feedback across time. BioRxiv. 10.64898/2026.02.25.707903

Thorne, D. R. (2006). Throughput: A simple performance index with desirable characteristics. Behavior Research Methods, 38(4), 569–573. 10.3758/BF03193886

Van Hulle, L., Juravle, G., Spence, C., Crombez, G., & Van Damme, S. (2013). Attention modulates sensory suppression during back movements. Consciousness and Cognition, 22(2), 420–429. 10.1016/j.concog.2013.01.011

Vitello, M. P., Fritschi, M., & Ernst, M. O. (2012). Active Movement Reduces the Tactile Discrimination Performance. In A. Peer & C. D. Giachritsis (Eds.), Immersive Multimodal Interactive Presence (pp. 7–34). Springer London. 10.1007/978-1-4471-2754-3_2

Voudouris, D., & Fiehler, K. (2017). Enhancement and suppression of tactile signals during reaching. Journal of Experimental Psychology: Human Perception and Performance, 43(6), 1238–1248. 10.1037/xhp0000373

Voudouris, D., & Fiehler, K. (2021). Dynamic temporal modulation of somatosensory processing during reaching. Scientific Reports, 11(1), 1928. 10.1038/s41598-021-81156-0

Wickham, H. & RStudio. (2023). tidyverse: Easily install and load the “Tidyverse” (Version 2.0.0) [Computer software]. https://cran.r-project.org/web/packages/tidyverse/index.html

Williams, S. R., Shenasa, J., & Chapman, C. E. (1998). Time course and magnitude of movement-related gating of tactile detection in humans. I. Importance of stimulus location. Journal of Neurophysiology, 79(2), 947–963. 10.1152/jn.1998.79.2.947

Wilson, M. R., Poolton, J. M., Malhotra, N., Ngo, K., Bright, E., & Masters, R. S. W. (2011). Development and validation of a surgical workload measure: The Surgery Task Load Index (SURG-TLX). World Journal of Surgery, 35(9), 1961–1969. 10.1007/s00268-011-1141-4

Zhai, S. (2004). Characterizing computer input with Fitts’ law parameters—The information and non-information aspects of pointing. International Journal of Human-Computer Studies, 61(6), 791–809. 10.1016/j.ijhcs.2004.09.006

