## Supplementary Analyses for "Vibrotactile stimulation on the upper back requires stronger vibration during goal-directed reaching but does not impede reaching performance"

This R markdown report contains 4 motor performance analyses not included in the main manuscript:

- Analysis 1: Peak Velocity

- Analysis 2: Time to Peak Velocity

- Analysis 3: Average Velocity

- Analysis 4: Acceleration

Analysis 1, 2, and 4 have two versions, a and b, due to an extreme outlier participant. Version a runs the analysis with the full dataset, and version b runs the analysis with the outlier participant’s data filtered out.

### Analysis 1a: Peak Velocity by Task, Target Size, and Distance

Peak velocity was defined as the maximum velocity attained between any two samples during the movement phase of each trial. Differences were tested with a three-way within-subjects ANOVA (task x target size x distance).

#### ANOVA summary

The ANOVA summary shows an effect of distance. Task and target size did not affect peak velocity, and there were no interactions.

```
summary(model)
##
## Univariate Type III Repeated-Measures ANOVA Assuming Sphericity
##
##                            Sum Sq num Df Error SS den Df  F value    Pr(>F)
## (Intercept)               1425449      1   272425     28 146.5084 1.215e-12 ***
## Task                        12968      2   138573     56   2.6204   0.08168 .
## Target_Size                     8      1      202     28   1.0868   0.30610
## Distance                     1656      1     6163     28   7.5218   0.01051 *
## Task:Target_Size               37      2      690     56   1.5071   0.23039
## Task:Distance                  76      2     4679     56   0.4564   0.63590
## Target_Size:Distance          101      1     1795     28   1.5817   0.21891
## Task:Target_Size:Distance      14      2      502     56   0.7715   0.46717
## ---
## Signif. codes:  0 '***' 0.001 '**' 0.01 '*' 0.05 '.' 0.1 ' ' 1
##
##
## Mauchly Tests for Sphericity
##
##                           Test statistic   p-value
## Task                             0.09633 0.0000000
## Task:Target_Size                 0.75055 0.0207799
## Task:Distance                    0.07927 0.0000000
## Task:Target_Size:Distance        0.47448 0.0000426
##
##
## Greenhouse-Geisser and Huynh-Feldt Corrections
##  for Departure from Sphericity
##
##                            GG eps Pr(>F[GG])
## Task                      0.52530     0.1149
## Task:Target_Size          0.80035     0.2330
## Task:Distance             0.52064     0.5123
## Task:Target_Size:Distance 0.65551     0.4188
##
##                              HF eps Pr(>F[HF])
## Task                      0.5281649  0.1146468
## Task:Target_Size          0.8413202  0.2326516
## Task:Distance             0.5229646  0.5131778
## Task:Target_Size:Distance 0.6748082  0.4221141
```

#### Plot

The plot shows that peak velocity was faster for the far targets, when participants had to move further to reach the target. The individual data reveals, however, that one participant had a peak velocity that was much faster than any other participant. This was especially the case while this participant was wearing the motors on the upper back during the reach-and-detect task. This severe outlier may be distorting the ANOVA outcome. We therefore re-ran the ANOVA without this participant in section 2b, below.

print(p_main)

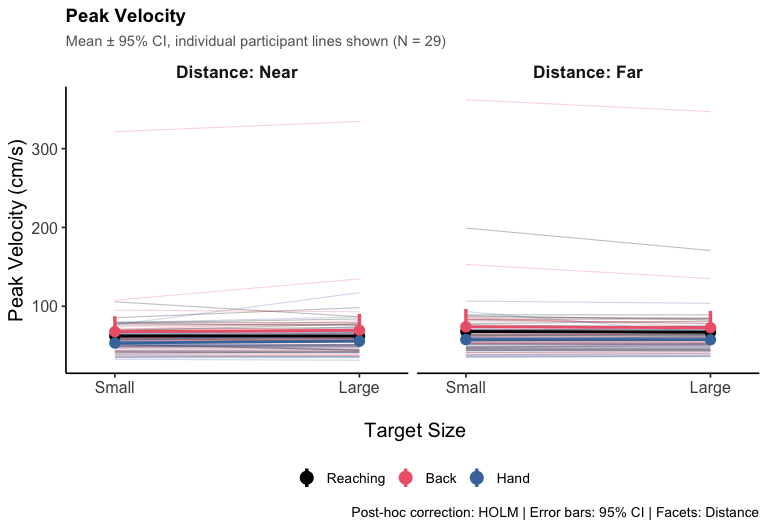

### Analysis 1b: Peak Velocity by Task, Target Size, and Distance (Filtered)

This variant re-runs peakVelocity.R with an outlier-exclusion range using customizable MIN_VAL/MAX_VAL lines, which are set to infinity by default in the original script.

We set the maximum value at 250 to exclude the outlier in the above data.

FILTER_MIN <- 0
FILTER_MAX <- 250

#### ANOVA summary (filtered)

The filter reduces our sample size from 29 to 28 participants.

The ANOVA summary on the filtered daya shows an effect of distance and task. Target size still did not affect peak velocity, and there were no interactions.

```
summary(model_filtered)
##
## Univariate Type III Repeated-Measures ANOVA Assuming Sphericity
##
##                            Sum Sq num Df Error SS den Df  F value    Pr(>F)
## (Intercept)               1184455      1    63739     27 501.7364 < 2.2e-16 ***
## Task                         3983      2    13775     54   7.8068  0.001052 **
## Target_Size                     3      1      173     27   0.4738  0.497136
## Distance                      752      1     1595     27  12.7383  0.001367 **
## Task:Target_Size                8      2      331     54   0.6439  0.529212
## Task:Distance                  44      2      726     54   1.6473  0.202097
## Target_Size:Distance           17      1      811     27   0.5514  0.464154
## Task:Target_Size:Distance      20      2      461     54   1.1722  0.317446
## ---
## Signif. codes:  0 '***' 0.001 '**' 0.01 '*' 0.05 '.' 0.1 ' ' 1
##
##
## Mauchly Tests for Sphericity
##
##                           Test statistic p-value
## Task                             0.73247 0.01747
## Task:Target_Size                 0.92935 0.38578
## Task:Distance                    0.39317 0.00001
## Task:Target_Size:Distance        0.41145 0.00001
##
##
## Greenhouse-Geisser and Huynh-Feldt Corrections
##  for Departure from Sphericity
##
##                            GG eps Pr(>F[GG])
## Task                      0.78894   0.002644 **
## Task:Target_Size          0.93401   0.519261
## Task:Distance             0.62234   0.210533
## Task:Target_Size:Distance 0.62951   0.300143
## ---
## Signif. codes:  0 '***' 0.001 '**' 0.01 '*' 0.05 '.' 0.1 ' ' 1
##
##                              HF eps  Pr(>F[HF])
## Task                      0.8296014 0.002212388
## Task:Target_Size          1.0008114 0.529211879
## Task:Distance             0.6377566 0.210405164
## Task:Target_Size:Distance 0.6459035 0.301321060
```

#### Plot (filtered)

With the filter, there is still one outlier participant relative to the average participant, but their values are not nearly so extreme as the filtered participant.

Peak velocity is still faster for the far targets than the near targets.

Peak velocity is notably slower when vibration was delivered to the hand during the reach-and-detect task, compared to when it was delivered to the upper back, or on the Reaching Task when there was no vibration. Perceiving or monitoring for vibration on the hand may cause people to limit the velocity of their movements.

print(p_filtered)

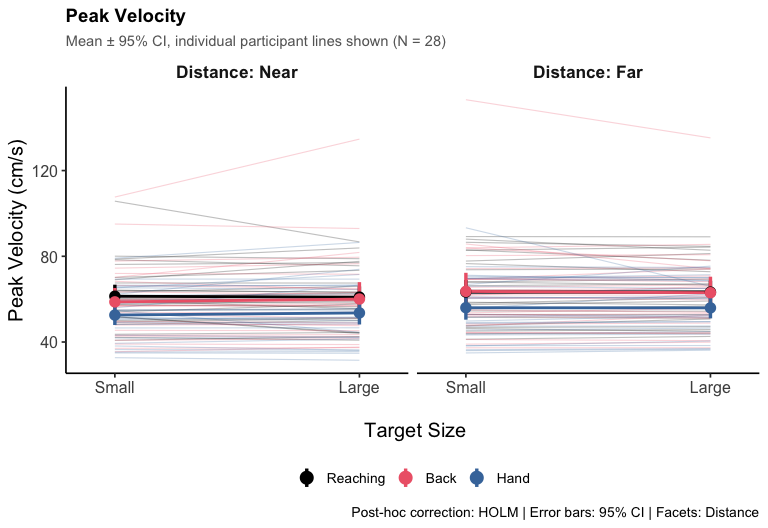

### Analysis 2a: Timing of Peak Velocity by Task, Target Size, and Distance

To calculate time to peak velocity, the duration of every movement was normalized to a value of 100, such that time of peak velocity represents a percentage of movement duration and is comparable for across both target distances, where MT is systematically different. A peak velocity time below 50% means that peak velocity was attained in the first half of the movement, while a time above 50% means it was attained in the second half of the movement. Differences were tested with a three-way within-subjects ANOVA (task x target size x distance).

#### ANOVA summary

The ANOVA summary shows effects of task, target size, and distance. There was also a target size x distance interaction.

```
summary(model)
##
## Univariate Type III Repeated-Measures ANOVA Assuming Sphericity
##
##                           Sum Sq num Df Error SS den Df   F value    Pr(>F)
## (Intercept)               442705      1   3387.0     28 3659.7635 < 2.2e-16 ***
## Task                         121      2    963.4     56    3.5071 0.0367286 *
## Target_Size                 2179      1    149.4     28  408.2762 < 2.2e-16 ***
## Distance                     832      1    175.1     28  132.9476 3.799e-12 ***
## Task:Target_Size               6      2    118.1     56    1.4732 0.2379316
## Task:Distance                 13      2    149.6     56    2.4149 0.0986335 .
## Target_Size:Distance          30      1     58.9     28   14.4750 0.0007078 ***
## Task:Target_Size:Distance      7      2     99.2     56    1.9792 0.1477276
## ---
## Signif. codes:  0 '***' 0.001 '**' 0.01 '*' 0.05 '.' 0.1 ' ' 1
##
##
## Mauchly Tests for Sphericity
##
##                           Test statistic p-value
## Task                             0.71809 0.01144
## Task:Target_Size                 0.92449 0.34648
## Task:Distance                    0.97600 0.72035
## Task:Target_Size:Distance        0.96675 0.63346
##
##
## Greenhouse-Geisser and Huynh-Feldt Corrections
##  for Departure from Sphericity
##
##                            GG eps Pr(>F[GG])
## Task                      0.78008    0.04941 *
## Task:Target_Size          0.92979    0.23882
## Task:Distance             0.97656    0.10007
## Task:Target_Size:Distance 0.96782    0.14938
## ---
## Signif. codes:  0 '***' 0.001 '**' 0.01 '*' 0.05 '.' 0.1 ' ' 1
##
##                              HF eps Pr(>F[HF])
## Task                      0.8177980 0.04695864
## Task:Target_Size          0.9932484 0.23802624
## Task:Distance             1.0488847 0.09863350
## Task:Target_Size:Distance 1.0384556 0.14772759
```

#### Plot

The plot shows that peak velocity was reached later in the trial for the reach-and-detect task, regardless of motor location, compared to the Reaching task without vibration. Perceiving or monitoring for vibration in either location may cause people to accelerate more slowly as they attend to the vibration. This is tested in Analysis 5 below.

Peak velocity was aslo reached earlier for small targets and for far distances. The greater precision required on these trials may result in participants taking longer to perform corrective movements near the end of the reach.

There is one outlier participant in this dataset, which is the same outlier as in the Peak Velocity analysis. To see if this has any effects on the results, we removed this participant and re-ran the ANOVA in section 3b, below.

print(p_main)

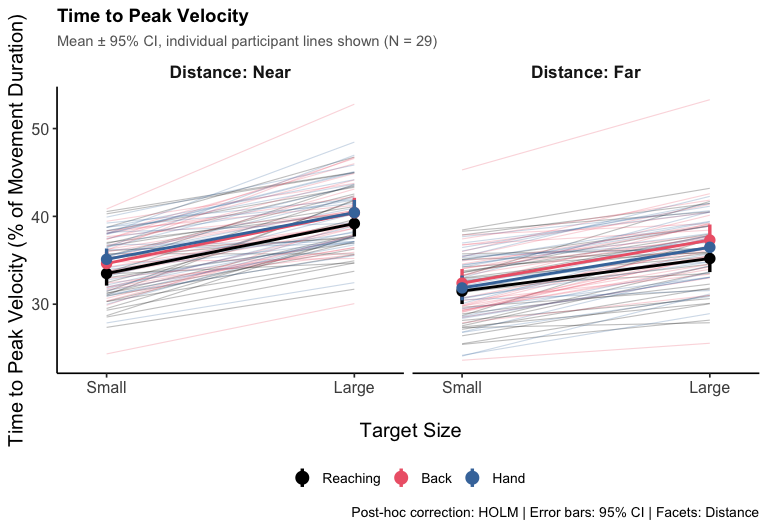

### Analysis 2b: Time to Peak Velocity by Task, Target Size, and Distance (Filtered)

This variant re-runs PV_timing.R with an outlier-exclusion range using customizable MIN_VAL/MAX_VAL lines, which are set to infinity by default in the original script.

We set the maximum value at 50 to exclude the same outlier participant as in the Peak Velocity analysis. This participant was the only participant to have a Peak Velocity time in the second half of movement for any condition.

FILTER_MIN <- 0
FILTER_MAX <- 50

#### ANOVA summary (filtered)

The filter reduces our sample size from 29 to 28 participants.

Filtering the outlier changes the outcome of the task comparison, if Greenhouse-Geiser sphericity correction is applied. Task differences are no longer significant. Task, target size, and distance remain significant. There remains a target size x distance interaction.

```
summary(model_filtered)
##
## Univariate Type III Repeated-Measures ANOVA Assuming Sphericity
##
##                           Sum Sq num Df Error SS den Df   F value    Pr(>F)
## (Intercept)               421788      1  2841.77     27 4007.4626 < 2.2e-16 ***
## Task                          94      2   791.02     54    3.2172  0.047857 *
## Target_Size                 2027      1   128.71     27  425.1722 < 2.2e-16 ***
## Distance                     852      1   153.61     27  149.8237 1.575e-12 ***
## Task:Target_Size               4      2   109.85     54    0.9029  0.411421
## Task:Distance                  7      2   128.91     54    1.5158  0.228821
## Target_Size:Distance          24      1    50.47     27   12.7551  0.001359 **
## Task:Target_Size:Distance      8      2    97.24     54    2.2315  0.117172
## ---
## Signif. codes:  0 '***' 0.001 '**' 0.01 '*' 0.05 '.' 0.1 ' ' 1
##
##
## Mauchly Tests for Sphericity
##
##                           Test statistic p-value
## Task                             0.74851 0.02315
## Task:Target_Size                 0.91080 0.29681
## Task:Distance                    0.99919 0.98955
## Task:Target_Size:Distance        0.95813 0.57350
##
##
## Greenhouse-Geisser and Huynh-Feldt Corrections
##  for Departure from Sphericity
##
##                            GG eps Pr(>F[GG])
## Task                      0.79905    0.06024 .
## Task:Target_Size          0.91810    0.40434
## Task:Distance             0.99919    0.22884
## Task:Target_Size:Distance 0.95981    0.11955
## ---
## Signif. codes:  0 '***' 0.001 '**' 0.01 '*' 0.05 '.' 0.1 ' ' 1
##
##                              HF eps Pr(>F[HF])
## Task                      0.8414092 0.05739571
## Task:Target_Size          0.9818422 0.40991270
## Task:Distance             1.0790265 0.22882098
## Task:Target_Size:Distance 1.0316760 0.11717193
```

#### Plot (filtered)

Removing the outlier participant does not notably change the pattern of the data.

print(p_filtered)

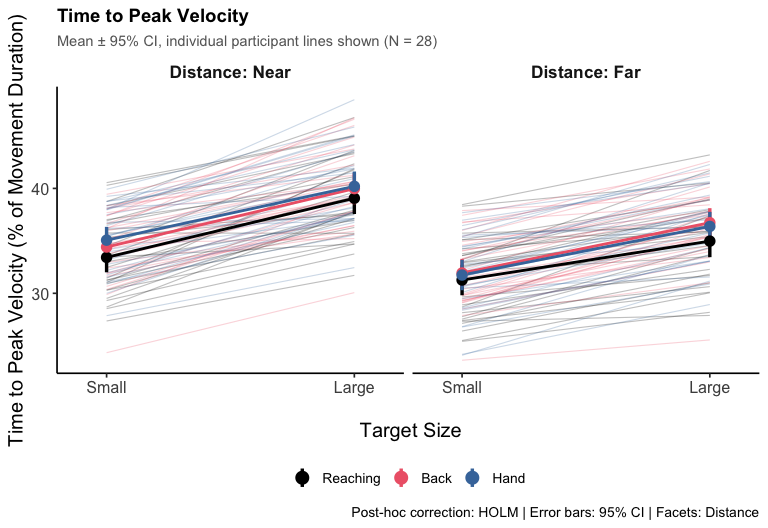

### Analysis 3: Average Velocity by Task, Target Size, and Distance

Average velocity was defined as the path length divided by movement time on each trial. Differences were tested with a three-way within-subjects ANOVA (task x target size x distance).

#### ANOVA summary

The ANOVA summary shows an effect of task, target size, and distance. There is also an interaction between task and target size.

```
summary(model)
##
## Univariate Type III Repeated-Measures ANOVA Assuming Sphericity
##
##                           Sum Sq num Df Error SS den Df   F value    Pr(>F)
## (Intercept)               254710      1   6958.7     28 1024.8818 < 2.2e-16 ***
## Task                         397      2    861.4     56   12.9116 2.448e-05 ***
## Target_Size                 1016      1    160.2     28  177.4938 1.216e-13 ***
## Distance                       6      1     37.5     28    4.7695   0.03750 *
## Task:Target_Size               4      2     33.8     56    3.5771   0.03451 *
## Task:Distance                  2      2     24.6     56    1.7207   0.18827
## Target_Size:Distance           0      1     11.3     28    0.7209   0.40306
## Task:Target_Size:Distance      1      2     20.1     56    0.9685   0.38591
## ---
## Signif. codes:  0 '***' 0.001 '**' 0.01 '*' 0.05 '.' 0.1 ' ' 1
##
##
## Mauchly Tests for Sphericity
##
##                           Test statistic p-value
## Task                             0.67264 0.00473
## Task:Target_Size                 0.99807 0.97427
## Task:Distance                    0.88412 0.18961
## Task:Target_Size:Distance        0.81782 0.06620
##
##
## Greenhouse-Geisser and Huynh-Feldt Corrections
##  for Departure from Sphericity
##
##                            GG eps Pr(>F[GG])
## Task                      0.75338  0.0001678 ***
## Task:Target_Size          0.99807  0.0346066 *
## Task:Distance             0.89615  0.1919629
## Task:Target_Size:Distance 0.84589  0.3742494
## ---
## Signif. codes:  0 '***' 0.001 '**' 0.01 '*' 0.05 '.' 0.1 ' ' 1
##
##                              HF eps   Pr(>F[HF])
## Task                      0.7869128 0.0001290117
## Task:Target_Size          1.0746161 0.0345136621
## Task:Distance             0.9534735 0.1899555326
## Task:Target_Size:Distance 0.8944325 0.3782078372
```

#### Plot

The plot shows that average velocity was fastest for the reaching task without vibration, slower when vibration was on the upper back, and slowest when the vibration was on the hand. This corresponds to the MT and peak velocity data, showing slower movements when perceiving and monitoring for vibration on the hand.

Average velocity was faster for the large targets, but also the far distances, albeit very slightly. In terms of average velocity, participants may have traded off precision to obtain a faster speed when they were required to reach further. This result could also be because longer movements contain proportionately less time spent accelerating and decelerating.

print(p_main)

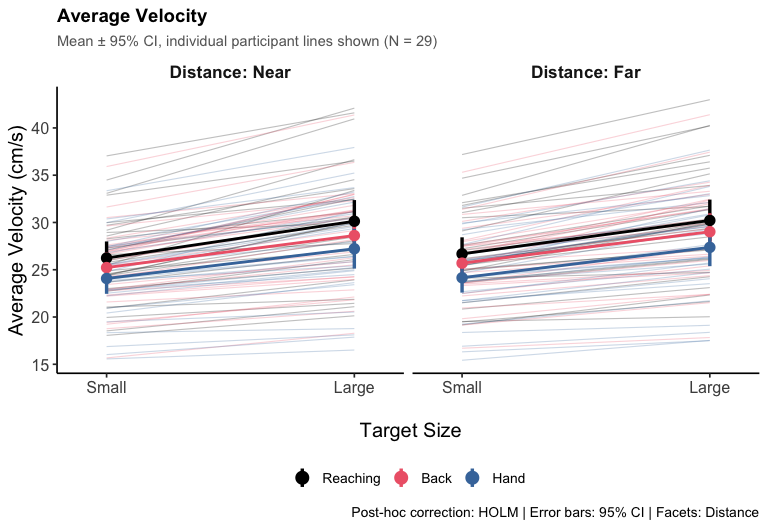

### Analysis 4a: Acceleration by Task, Target Size, and Distance

Acceleration was defined as peak velocity divided by time to peak velocity. Differences were tested with a three-way within-subjects ANOVA (task x target size x distance).

#### ANOVA summary

The ANOVA summary shows an effect of task and distance.

```
summary(model)
##
## Univariate Type III Repeated-Measures ANOVA Assuming Sphericity
##
##                             Sum Sq num Df Error SS den Df  F value    Pr(>F)
## (Intercept)               26923057      1  4602486     28 163.7910 3.213e-13
## Task                        346835      2  1197749     56   8.1080 0.0008082
## Target_Size                    611      1     4600     28   3.7202 0.0639529
## Distance                     23611      1    32649     28  20.2486 0.0001087
## Task:Target_Size               785      2    16968     56   1.2950 0.2819834
## Task:Distance                   70      2    46863     56   0.0415 0.9593393
## Target_Size:Distance            72      1     9331     28   0.2173 0.6447127
## Task:Target_Size:Distance      984      2    20164     56   1.3670 0.2632459
##
## (Intercept)               ***
## Task                      ***
## Target_Size               .
## Distance                  ***
## Task:Target_Size
## Task:Distance
## Target_Size:Distance
## Task:Target_Size:Distance
## ---
## Signif. codes:  0 '***' 0.001 '**' 0.01 '*' 0.05 '.' 0.1 ' ' 1
##
##
## Mauchly Tests for Sphericity
##
##                           Test statistic   p-value
## Task                             0.74036 0.0172765
## Task:Target_Size                 0.60706 0.0011847
## Task:Distance                    0.45037 0.0000210
## Task:Target_Size:Distance        0.39549 0.0000036
##
##
## Greenhouse-Geisser and Huynh-Feldt Corrections
##  for Departure from Sphericity
##
##                            GG eps Pr(>F[GG])
## Task                      0.79388   0.002085 **
## Task:Target_Size          0.71791   0.276466
## Task:Distance             0.64532   0.894243
## Task:Target_Size:Distance 0.62324   0.258199
## ---
## Signif. codes:  0 '***' 0.001 '**' 0.01 '*' 0.05 '.' 0.1 ' ' 1
##
##                              HF eps  Pr(>F[HF])
## Task                      0.8337947 0.001734429
## Task:Target_Size          0.7460889 0.277370465
## Task:Distance             0.6632183 0.899388546
## Task:Target_Size:Distance 0.6382002 0.258712699
```

#### Plot

The plot shows that acceleration was slower when the vibration was on the hand, compared to when it was on the upper back or on the reaching task without vibration. This corresponds to the MT and peak velocity data, showing slower movements when perceiving and monitoring for vibration on the hand. The vibrations in this study were delivered before movement onset, at movement onset, or 500ms after movement onset; in other words, during the earlier part of the trial, rather than at the end of the trial when participants have reached the targets. Overall MT may be slower when the vibration is delivered to the hand because they are accelerating more slowly as they process or monitor for the vibration.

Corresponding to the average velocity data, acceleration was faster for far targets. Trials with far targets would take longer to complete than trials with near targets, all else being equal, so participants may be prioritizing speed at the start of these trials, knowing they can correct for precision once they approach the target.

There is one outlier participant with notably fast acceleration, relative to other participants when the upper back was vibrated. This is the same participant with outlier values for peak velocity and time to peak velocity. We therefore re-ran the acceleration analysis without this participant in section 5b, below.

print(p_main)

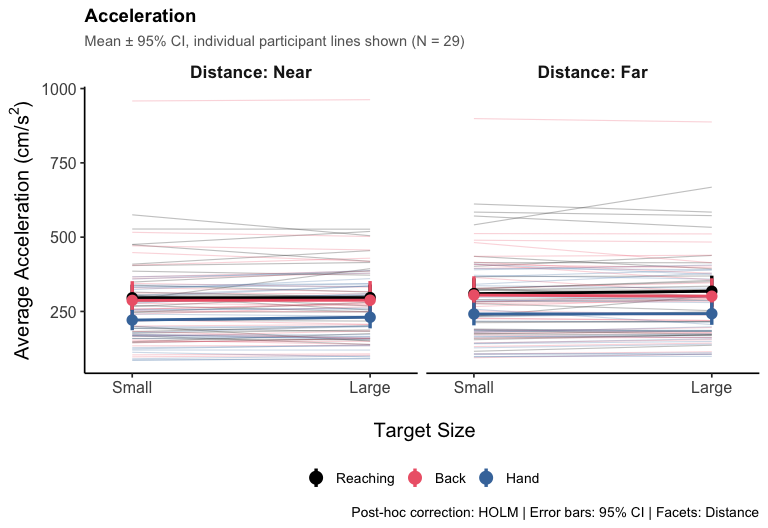

### Analysis 4b: Acceleration by Task, Target Size, and Distance (Filtered)

This variant re-runs acceleration.R with an outlier-exclusion range using customizable MIN_VAL/MAX_VAL lines, which are set to infinity by default in the original script.

We set the maximum value at 900 to exclude the outlier in the above data.

FILTER_MIN <- 0
FILTER_MAX <- 900

#### ANOVA summary (filtered)

The filter reduces our sample size from 29 to 28 participants.

The ANOVA summary on the filtered daya shows an effect of distance and task. Target size still did not affect peak velocity, and there were no interactions.

```
summary(model_filtered)
##
## Univariate Type III Repeated-Measures ANOVA Assuming Sphericity
##
##                             Sum Sq num Df Error SS den Df  F value    Pr(>F)
## (Intercept)               23865218      1  3283158     27 196.2626 6.646e-14
## Task                        270853      2   606014     54  12.0674 4.654e-05
## Target_Size                    468      1     4395     27   2.8753 0.1014536
## Distance                     19448      1    28794     27  18.2362 0.0002157
## Task:Target_Size               624      2    16665     54   1.0117 0.3703810
## Task:Distance                  533      2    19216     54   0.7489 0.4777280
## Target_Size:Distance             7      1     8388     27   0.0228 0.8812111
## Task:Target_Size:Distance     1295      2    19353     54   1.8065 0.1740182
##
## (Intercept)               ***
## Task                      ***
## Target_Size
## Distance                  ***
## Task:Target_Size
## Task:Distance
## Target_Size:Distance
## Task:Target_Size:Distance
## ---
## Signif. codes:  0 '***' 0.001 '**' 0.01 '*' 0.05 '.' 0.1 ' ' 1
##
##
## Mauchly Tests for Sphericity
##
##                           Test statistic p-value
## Task                             0.99711 0.96302
## Task:Target_Size                 0.56992 0.00067
## Task:Distance                    0.81042 0.06505
## Task:Target_Size:Distance        0.40755 0.00001
##
##
## Greenhouse-Geisser and Huynh-Feldt Corrections
##  for Departure from Sphericity
##
##                            GG eps Pr(>F[GG])
## Task                      0.99711  4.751e-05 ***
## Task:Target_Size          0.69926     0.3477
## Task:Distance             0.84063     0.4570
## Task:Target_Size:Distance 0.62796     0.1878
## ---
## Signif. codes:  0 '***' 0.001 '**' 0.01 '*' 0.05 '.' 0.1 ' ' 1
##
##                              HF eps   Pr(>F[HF])
## Task                      1.0765189 0.0000465398
## Task:Target_Size          0.7257122 0.3502270585
## Task:Distance             0.8901600 0.4638443878
## Task:Target_Size:Distance 0.6441459 0.1873627521
```

#### Plot (filtered)

With the outlier participant removed, acceleration remains slower when the hand is vibrated, compared to when the upper back is vibrated or when there is no vibration in the reaching task.

Acceleration also remains faster for the far targets.

print(p_filtered)

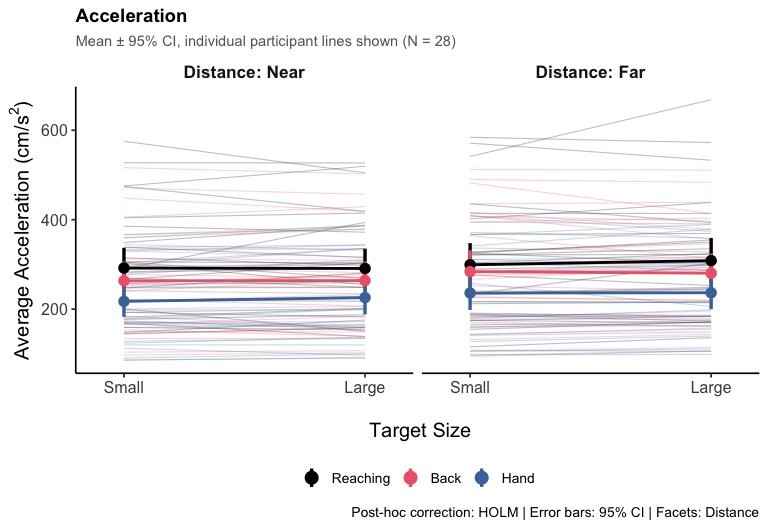
